# Gene network analysis identifies a central post-transcriptional regulator of cellular stress survival

**DOI:** 10.1101/212902

**Authors:** Matthew Z. Tien, Aretha Fiebig, Sean Crosson

**Affiliations:** Department of Biochemistry and Molecular Biology, University of Chicago, Chicago, IL. USA. 929 E. 57^th^ St. - GCIS W138 Chicago, IL 60637; Department of Microbiology, University of Chicago, Chicago, IL. USA

## Abstract

Cells adapt to shifts in their environment by remodeling transcription. Measuring changes in transcription at the genome scale is now routine, but defining the functional significance of individual genes within large gene expression datasets remains a major challenge. We applied a network-based algorithm to interrogate publicly available transcription data to predict genes that serve major functional roles in *Caulobacter crescentus* stress survival. This approach identified GsrN, a conserved small RNA that is directly controlled by the general stress sigma factor, σ^τ^, and functions as a potent post-transcriptional regulator of survival under multiple stress conditions. GsrN expression is both necessary and sufficient to protect cells from hydrogen peroxide, where it functions by base pairing with the leader of *katG* mRNA and promoting catalase/peroxidase expression. We conclude that GsrN convenes a post-transcriptional layer of gene expression that serves a central functional role in stress physiology.

## INTRODUCTION

Organisms must control gene expression to maintain homeostasis. A common mode of gene regulation in bacteria involves activation of alternative sigma factors (σ), which redirect RNA polymerase to transcribe genes required for adaptation to particular environmental conditions. Alphaproteobacteria utilize an extracytoplasmic function (ECF) σ factor to initiate a gene expression program known as the general stress response (GSR) (Figure 1A). The GSR activates transcription of dozens of genes, which mitigates the detrimental effects of environmental stressors and influences the infection biology of alphaproteobacterial pathogens (reviewed in (Fiebig et al., 2015; Francez-Charlot et al., 2015)). The molecular mechanisms by which genes in the GSR regulon enable growth and survival across a chemically- and physically-distinct spectrum of conditions are largely uncharacterized. Defining the functional role(s) of individual genes contained within complex environmental response regulons, such as the GSR, remains a major challenge in microbial genomics.

**Figure 1.**
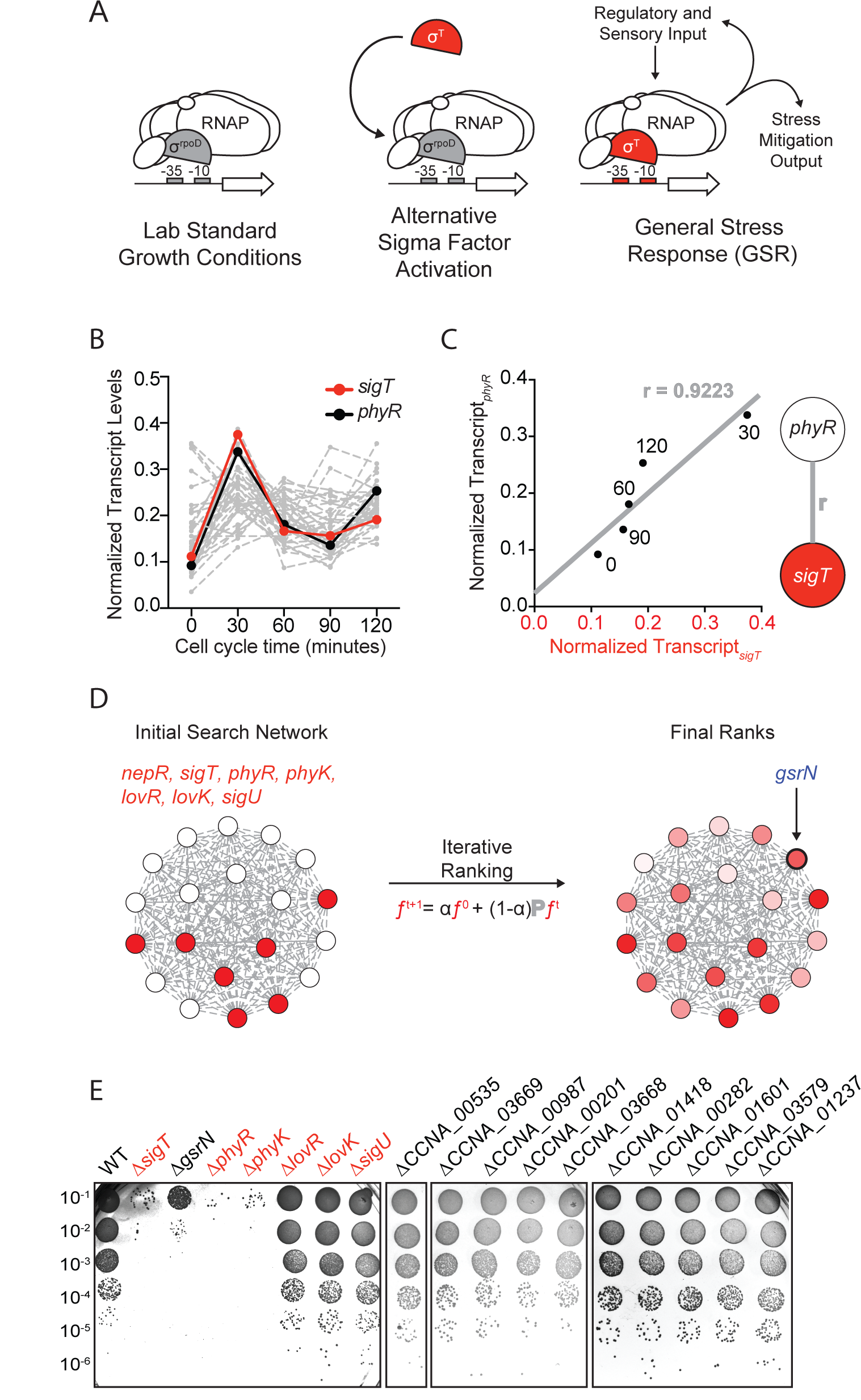
Iterative rank analysis of gene expression data identifies *gsrN*, a small RNA that confers resistance to hydrogen peroxide. (A) Activation of general stress response (GSR) sigma factor, σ^T^, promotes transcription of genes that mitigate the effects of environmental stress and genes that regulate σ^T^ activity. (B) Normalized transcript levels from (Fang et al., 2013) of known GSR regulated genes are plotted as a function of cell cycle time. The core GSR regulators, *sigT* and *phyR*, are highlighted in red and black respectively. Data plotted from Figure 1- source data 1. (C) *sigT* and *phyR* transcript levels are correlated as a function of cell cycle progression, Pearson’s correlation coefficient *r* = 0.92. (D) An initial correlation-weighted network was seeded with experimentally-defined GSR regulatory genes (red, value=1) (left). Final ranks were calculated using the stable solution of the iterative ranking algorithm (right). Red intensity scales with the final rank weights (Figure 1- source data 2). A gene encoding a small RNA, *gsrN*, was a top hit on the ranked list. (E) Colony forming units (CFU) in dilution series (10^-1^ to 10^-6^) of wild-type and mutant *Caulobacter* strains after 0.2 mM hydrogen peroxide treatment for 1 hour. Red denotes core GSR regulatory genes. Black denotes known σ^T^–regulated genes, listed by GenBank locus ID.

In the alphaproteobacterium *Caulobacter crescentus*, GSR mutant strains have survival defects under multiple conditions including hyperosmotic and hydrogen peroxide stresses (Alvarez-Martinez et al., 2007; Foreman et al., 2012). However, the majority of genes regulated at the transcriptional level by the *Caulobacter* GSR sigma factor, σ^T^, have no annotated function or no clear role in stress physiology. While studies of transcription can provide understanding of stress responses, this approach may miss functionally important processes that are regulated at the post-transcriptional level, such as those controlled by small RNAs (sRNAs). Roles for sRNAs in bacterial stress response systems are well described (Wagner and Romby, 2015), but remain unexplored in the alphaproteobacterial GSR.

sRNAs typically function as repressors, though the regulatory roles and mechanisms of action of these molecules are diverse: sRNAs can control gene expression by protein sequestration, modulation of mRNA stability, transcription termination, or promotion of translation (Wagner and Romby, 2015). The system properties of environmental response networks are often influenced by sRNAs, which can affect the dynamics of gene expression via feedback (Beisel and Storz, 2011; Mank et al., 2013; Nitzan et al., 2015; Shimoni et al., 2007) or buffer response systems against transcriptional noise (Arbel-Goren et al., 2013; Golding et al., 2005; Levine and Hwa, 2008; Mehta et al., 2008). However, the phenotypic consequences of deleting sRNA genes are typically subtle and uncovering phenotypes often requires cultivation under particular conditions. Thus, reverse genetic approaches to define functions of uncharacterized sRNAs have proven challenging.

We applied a rank-based network analysis approach to predict the most functionally significant genes in the *Caulobacter* GSR regulon. This analysis led to the prediction that a sRNA, which we name GsrN, is a major genetic determinant of growth and survival under stress. We validated this prediction, demonstrating that *gsrN* is under direct control of σ^T^ and functions as a potent post-transcriptional regulator of survival across distinct conditions including hydrogen peroxide stress and hyperosmotic shock. We developed a novel forward biochemical approach to identify direct molecular targets of GsrN and discovered that peroxide stress survival is mediated through an interaction between GsrN and the 5’ leader sequence of *katG*, which activates KatG catalase/peroxidase expression. This post-transcriptional connection between σ^T^ and *katG*, a major determinant of peroxide stress and stationary phase survival (Italiani et al., 2011; Steinman et al., 1997), explains the peroxide sensitivity phenotype of *Caulobacter* strains lacking a GSR system.

Finally, we demonstrate that RNA processing and sRNA-mRNA target interactions shape the pool of functional GsrN in the cell, and that changes in GsrN expression enhance expression of some proteins while inhibiting others. The broad regulatory capabilities of GsrN are reflected in the fact that a *gsrN* deletion strain has survival defects across chemically- and physically-distinct stress conditions, and support a model in which the GSR initiates layered transcriptional and post-transcriptional regulatory responses to ensure environmental stress survival.

## RESULTS

### Iterative rank analysis of gene expression data identifies a small RNA regulator of stress survival

We applied a network-based analytical approach to interrogate published transcriptomic datasets (Fang et al., 2013) and predict new functional genetic components of the *Caulobacter* GSR system. We organized expression data for over 4000 genes (Figure 1B and Figure 1- source data 1) to create a weighted network. In our basic network construction, each gene in the genome was represented as a node and each node was linked to every other node by a correlation coefficient that quantified the strength of co-expression across all datasets (Figure 1C). Within this undirected graph, we aimed to uncover a GSR clique and thus more explicitly define the core functional components of the GSR regulon.

To identify uncharacterized genes that are strongly associated with the GSR, we utilized an iterative ranking approach related to the well-known PageRank algorithm (Brin and Page, 1998). We defined the “input” set as the experimentally-defined regulators of σ^T^ (Figure 1D), optimized parameters through a systematic self-predictability approach (Figure 1- figure supplement 1A and Materials and methods-Iterative rank parameter tuning), and applied iterative ranking to compute a ranked list of genes with strong associations to the input set (Figure 1- source data 2). We narrowed our ranked list by performing a promoter motif search to predict direct targets of σ^T^. A gene encoding an sRNA with a consensus σ^T^ binding site in its promoter, *ccna_R0081* (Landt et al., 2008), was a top hit in our rank list. We hereafter refer to this gene as *gsrN* (<u>g</u>eneral <u>s</u>tress <u>r</u>esponse non-coding <u>R</u>NA).

To test whether *gsrN* transcription requires the GSR sigma factor, σ^T^, we generated a transcriptional reporter by fusing the *gsrN* promoter to *lacZ (P_gsrN_lacZ).* Transcription from P*_gsrN_* required *sigT* (Figure 1- figure supplements 2A,C), validating *gsrN* as a bona fide member of the GSR regulon. To determine whether *gsrN* is a feedback regulator of GSR transcription, we utilized a well-characterized P*_sigU_lacZ* reporter (Foreman et al., 2012). Transcription from P*_sigU_* required *sigT*and other GSR regulators *(phyR, phyK)*, but was unaffected by deletion or overexpression of *gsrN.* Additionally, transcription from P_sigU_ under known inducers of a GSR response through sucrose addition was not affected either (Figure 1- figure supplement 2D). We conclude *gsrN* is activated by σ^T^ but does not feedback to control GSR transcription.

We next tested whether *gsrN* plays a role in stress survival. We subjected strains lacking *gsrN* or the core GSR regulators, *sigT, phyR*, or *phyK*, to hydrogen peroxide, a known stress under which GSR regulatory mutants have a survival defect. Δ*sigT*, Δ*phyR*, and Δ*phyK* strains had a ≈4-log decrease in cell survival relative to wild type after exposure to hydrogen peroxide, as previously reported (Alvarez-Martinez et al., 2007; Foreman et al., 2012). Cells lacking *gsrN* (Δ*gsrN*) had a ≈3-log viability defect relative to wild type (Figures 1E and Figure 1- figure supplement 2B). Insertion of *gsrN* with its native promoter at the ectopic *vanA* locus fully complemented the peroxide survival defect of Δ*gsrN* (Figure 2A). These data provide evidence that *gsrN* is a major genetic contributor to cell survival upon peroxide exposure. We identified 10 additional genes that are strongly regulated by σ^T^ and generated strains harboring single, in-frame deletions of these genes. The functions of these 10 genes are unknown: 6 encode conserved hypothetical proteins; 2 encode predicted outer membrane proteins; 1 encodes a cold shock protein, and 1 encodes a ROS/MUCR transcription factor. None of these additional deletion strains were sensitive to hydrogen peroxide (Figures 1E and Figure 1- figure supplement 2B).

**Figure 2.**
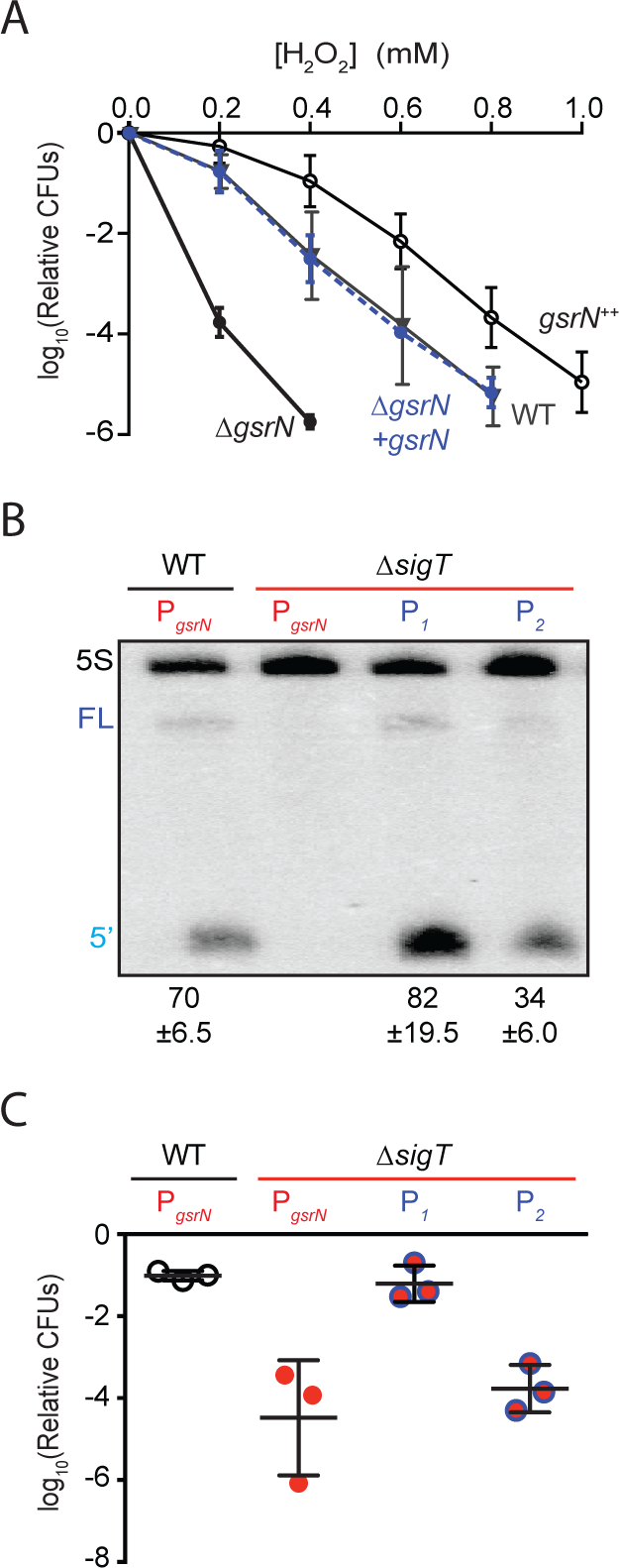
GsrN is necessary and sufficient for hydrogen peroxide stress survival. (A) *Caulobacter* wild type (WT), *gsrN* deletion (Δ*gsrN*), complementation (Δ*gsrN*+*gsrN*), and *gsrN* overexpression (4*gsrN*, *gsrN*^++^) strains were subjected to increasing concentrations of hydrogen peroxide for one hour and tittered on nutrient agar. Log_10_ relative CFU (peroxide treated/untreated) is plotted as a function of peroxide concentration. Δ*gsrN* and WT strains carried the empty plasmid (pMT552) as a control. Mean ± SD, n=3 independent replicates. (B) Northern blot of total RNA isolated from WT and Δ*sigT* strains expressing *gsrN* from its native promoter (P_*sigT*_) or from two constitutive σ^RpoD^ promoters (P_1_ or P_2_); probed with ^32^P-labeled oligonucleotides specific for GsrN and 5S rRNA as a loading control. Quantified values are mean ± SD of normalized signal, n=3 independent replicates. (C) Relative survival of strains in (B) treated with 0.2 mM hydrogen peroxide for 1 hour normalized as in (A). Mean ± SD from 3 independent experiments (points) is presented as bars.

### Expression of GsrN is sufficient for survival under peroxide stress

Results outlined above demonstrate that *gsrN* is necessary for hydrogen peroxide stress survival. To assess the effects of *gsrN* overexpression, we inserted constructs containing either one or three copies of *gsrN* under its native promoter into the *vanA* locus of wild-type and Δ*gsrN* strains (Figure 2 - figure supplement 1A). We measured GsrN expression directly in these strains by Northern blot (Figure 2 - figure supplement 1B) and tested their susceptibility to hydrogen peroxide (Figure 2A). Treatment with increasing concentrations of hydrogen peroxide revealed that strains overexpressing *gsrN* have a survival advantage compared to wild type. Measured levels of GsrN in the cell directly correlated *(r=0.92)* with cell survival, which provides evidence that the protective effect of *gsrN* under peroxide stress is dose dependent over the measured range (Figure 2 - figure supplement 1C).

To test sufficiency of *gsrN* to regulate cell survival under peroxide stress, we decoupled *gsrN* transcription from σ^T^. *gsrN* was constitutively expressed from promoters (P1 and P2) controlled by the primary sigma factor, RpoD, in a strain lacking *sigT* (Figure S2A). *gsrN* expression from P1 was 15% higher, and expression from P2 50% lower than *gsrN* expressed from its native σ^T^-dependent promoter (Figure 2B). Expression of *gsrN* from P1, but not P2, rescued the Δ*igT* peroxide survival defect (Figure 2C). We conclude that *gsrN* is the sole genetic determinant of hydrogen peroxide survival regulated downstream of σ^T^ under these conditions. Consistent with the dose dependent protection by GsrN, these data demonstrate that a threshold level of *gsrN* expression is required to protect the cell from hydrogen peroxide.

### GsrN is endonucleolytically processed into a more stable 5’ isoform

A notable feature of GsrN is the presence of two isoforms by Northern blot. Probes complementary to the 5’ portion of GsrN reveal full-length (≈100 nucleotide) and short (51 to 54 nucleotides) isoforms while probes complementary to the 3’ portion reveal mostly full-length GsrN (Figure 3A and Figure 3- figure supplement 1A). Two isoforms of GsrN are also evident in RNA-seq data (Figure 3- figure supplement 1B).

**Figure 3.**
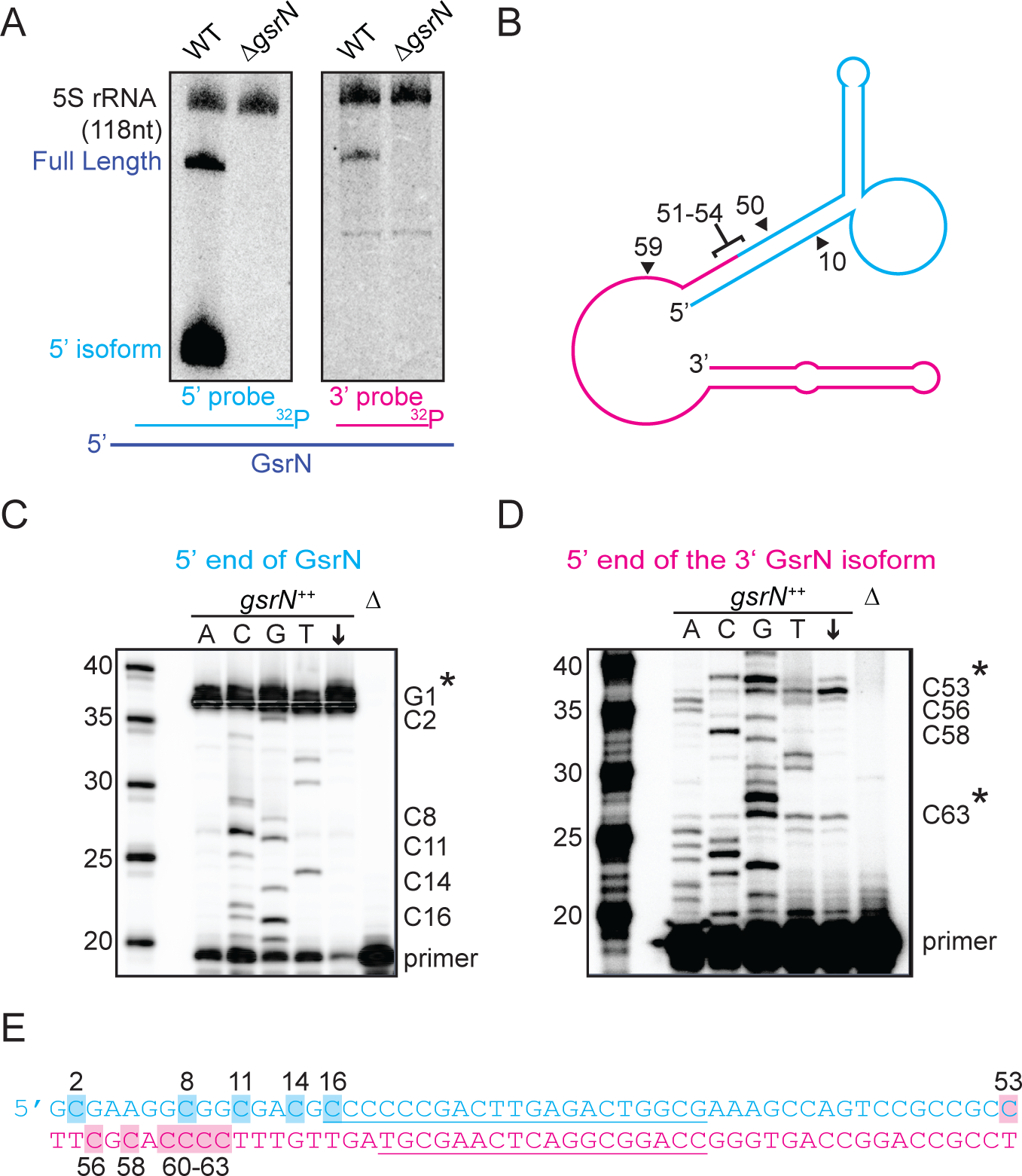
Full-length GsrN is endonucleolytically processed into a more stable 5’ isoform. (A) Northern blots of total RNA from wild-type and Δ*gsrN* cells hybridized with probes complementary to the 5’end (left) or 3’ end (right) of GsrN, and to 5S rRNA as a loading control. (B) Predicted secondary structure of full-length GsrN. Nucleotide positions labeled with arrows. Cyan indicates the 5’ end of GsrN determined by primer extension. Pink represents the 3’ end. (C) Primer extension from total RNA extracted from *gsrN^++^* and Δ*gsrN* (negative control) cultures (OD_660_ ≈ 1.0). Sequence was generated from a radiolabeled oligo anti-sense to the underlined cyan sequence in (E). Sanger sequencing control lanes A, C, G, and T mark the respective ddNTP added to that reaction to generate nucleotide specific stops. “C” labels on the right of the gel indicate mapped positions from the “G” lane. Arrow indicates lane without ddNTPs. Asterisk indicates positions of 5’ termini. (D) Primer extension from RNA samples as in (C). Sequence was extended from a radiolabeled oligo anti-sense to the underlined pink sequence in (E). (E) GsrN coding sequence. Cyan and pink indicate the predicted 5’ and 3’ isoforms, respectively. Primers binding sites used for primer extension in (C) and (D) are underlined. Highlighted C positions correspond to ddGTP stops in the “G” extensions.

The short isoform of *gsrN* could arise through two biological processes: alternative transcriptional termination or endonucleolytic processing of full-length GsrN. To test these two possibilities, we inhibited transcription with rifampicin, and monitored levels of both GsrN isoforms over time. Full-length GsrN decayed exponentially with a half-life of ^~^105 seconds (Figure 3- figure supplements 1C,D). The 5’ isoform increased in abundance for several minutes after treatment, concomitant with the decay of the full-length product. This observation is consistent with a model in which the 5’ isoform arises from the cleavage of the full-length product.

To identify potential endonucleolytic cleavage sites, we conducted primer extension assays, primer extension binding sites shown in (Figure 3E). Extension from an oligo complementary to the 5’ portion of GsrN confirmed the annotated transcriptional start site (Figure 3C). Extension from the 3’ portion identified two internal 5’ ends (Figure 3D). The positions of these internal 5’ ends are consistent with two small bands observed on Northern blots of high concentrations of total RNA hybridized with the 3’ probe (Figure 3- figure supplement 1A). The terminus around C53 corresponds to a potential endonucleolytic cleavage site that would generate the abundant stable 5’ isoform (Figure 3B).

### 5’ end of GsrN is necessary and sufficient for peroxide survival

To test the function of the 5’ portion of GsrN, we integrated a *gsrN* allele that contains only the first 58 nucleotides (Δ59-106), and lacks the transcriptional terminator (gsrNΔ3’) into the *vanA* locus (Figure 4A). This short *gsrN* allele complemented the Δ*gsrN* peroxide survival defect (Figure 4B). The *gsrNΔ3’* allele produced a 5’ isoform that was comparable in size and concentration to the wild-type 5’ *gsrN* isoform. Since the transcriptional terminator of *gsrN* was removed, we also observed a run-on ^~^200nt transcript from *gsrNΔ3’* (Figure 4C).

**Figure 4.**
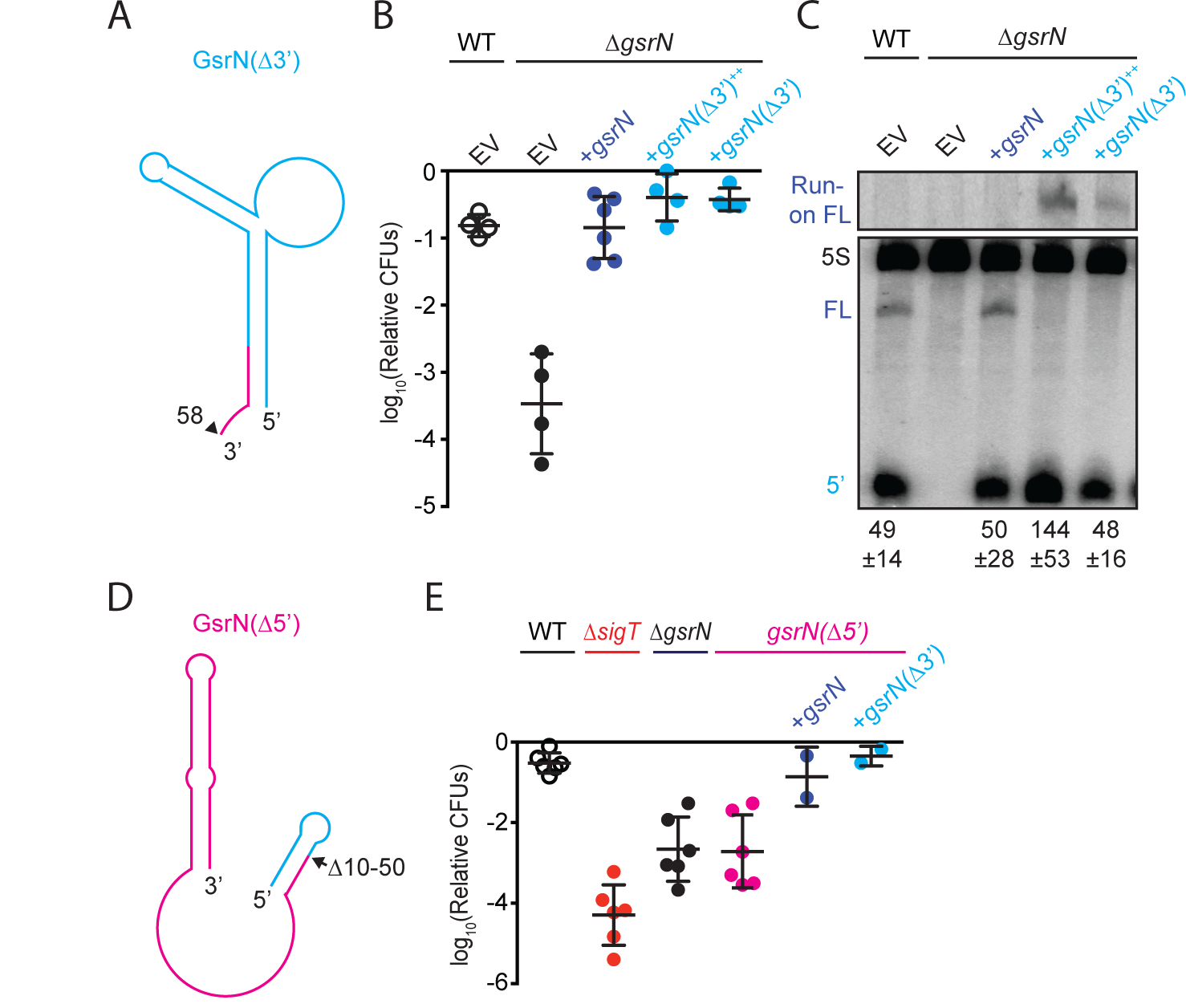
5’ end of GsrN is necessary and sufficient for peroxide survival. (A) Schematic diagram of GsrN(Δ3’), which lacks nucleotides 59-106. (B) Relative survival of strains treated with 0.2 mM hydrogen peroxide for 1 hour. WT and Δ*gsrN* strains carry empty plasmids (EV), plasmids harboring full-length *gsrN*, *gsrN*(Δ3’), or multiple copies of *gsrN*(Δ3’) (labeled *gsrN*(Δ3’)^++^). Bars represent mean ± SD from 4 independent experiments (points). (C) Northern blot of total RNA from strains in panel 3D harvested during exponential growth phase. Blots were hybridized with probes complementary to the 5’ end of GsrN and 5S rRNA. Mean ± SD of total GsrN signal from 3 independent samples. (D) Schematic diagram of GsrN(Δ5’), which lacks nucleotides 10-50. (E) Relative survival of strains treated with 0.2 mM hydrogen peroxide for 1 hour. Genetic backgrounds are indicated above the line; the GsrN(Δ5’) strain was complemented with either *gsrN* (dark blue) or GsrN(Δ5’) (cyan). Bars represent mean ± SD from several independent experiments (points).

To test the necessity of the 5’ portion of GsrN in peroxide stress survival, we deleted nucleotides 10 to 50 of *gsrN* at its native locus (Figure 4D). The *gsrNΔ5’* strain had a peroxide viability defect that was equivalent to *ΔgsrN.* Ectopic expression of either full-length *gsrN* or *gsrNΔ3’* in the *gsrNΔ5’* strain complemented its peroxide survival defect (Figure 4E).

### Several RNAs, including *katG* mRNA, co-purify with GsrN

We developed a forward biochemical approach to identify molecular partners of GsrN. The *<u>P</u>seudomonas* <u>p</u>hage<u>7</u> (PP7) genome contains hairpin (PP7hp) aptamers that bind to PP7 coat protein (PP7cp) with nanomolar affinity (Lim and Peabody, 2002). We inserted the PP7hp aptamer into multiple sites of *gsrN* with the goal of purifying GsrN with its interacting partners from *Caulobacter* lysates by affinity chromatography (Figure 5A), similar to an approach used by (Hogg and Collins, 2007; Said et al., 2009). PP7hp insertions at the 5’ end of *gsrN* and at several internal nucleotide positions (37, 54, 59, 67, and 93nt) were functionally assessed (Figure 5- figure supplement 1A). GsrN-PP7hp alleles tagged at the 5’ end or at nucleotide positions 54 or 59 did not complement the Δ*gsrN* peroxide survival defect (Figure 5- figure supplement 1B). These alleles yielded lower steady-state levels of 5’ isoform compared to wild type (Figure 5- figure supplements 1C,D). GsrN-PP7hp alleles with insertions at nucleotides 37, 67, and 93 restored peroxide resistance to Δ*gsrN* and produced more 5’ isoform than non-complementing GsrN-PP7 constructs (Figure 5- figure supplement 1).

**Figure 5.**
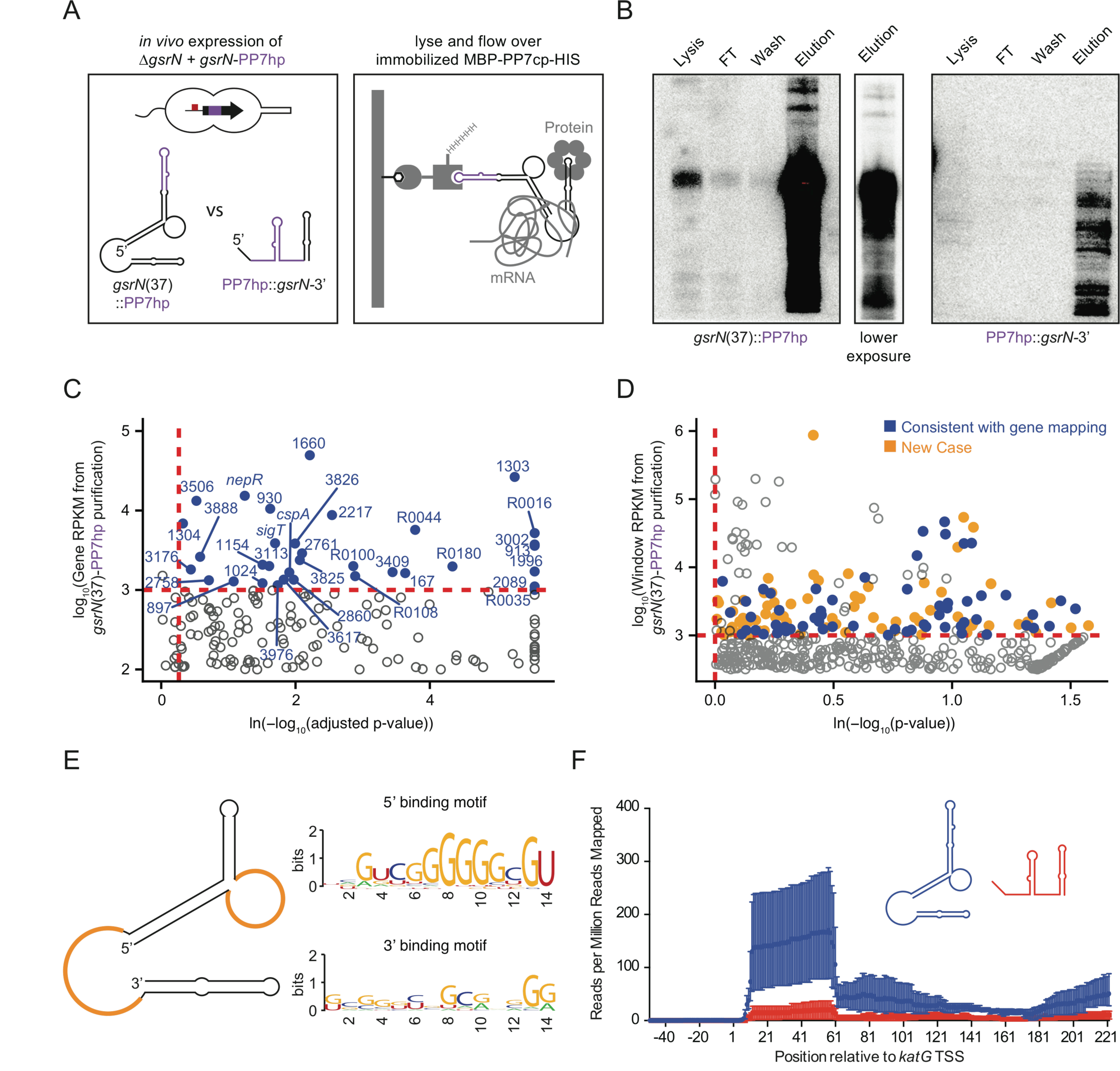
GsrN co-purifies with multiple RNAs, including catalase/peroxidase *katG* mRNA. (A) GsrN-target co-purification strategy. GsrN(black)-PP7hp(purple) fusions were expressed in a Δ*gsrN* background. PP7 RNA hairpin (PP7hp) inserted at nucleotide 37 (*gsrN*(37)⸬PP7hp) was used as the bait. PP7hp fused to the 3’ hairpin of *gsrN* (PP7hp⸬*gsrN*-3’)served as a negative control. Stationary phase cultures expressing these constructs were lysed and immediately flowed over an amylose resin column containing immobilized PP7hp binding protein (MBP-PP7cp-His). (B) GsrN-PP7hp purification from strains bearing *gsrN*(37)⸬PP7hp (left) and PP7hp⸬*gsrN*-3’ (right) was monitored by Northern Blot with probes complementary to 5’ end of GsrN and PP7hp, respectively. Lysate, flow through (FT), buffer wash, and elution fractions are blotted. Approximately 1μg RNA was loaded per lane, except for buffer wash (insufficient amount of total RNA). (C) Annotation-based analysis of transcripts that co-purify with *gsrN*(37)⸬PP7hp (Figure 5-source data 1). Log_10_ reads per kilobase per million reads (RPKM) is plotted against the ln(-log_10_(false discovery rate corrected p-value)). Dashed red lines mark the enrichment co-purification thresholds. Genes enriched in the gsrN(37)⸬PP7hp purification compared to PP7hp⸬gsrN-3’ are blue; labels correspond to gene names or *C. crescentus* strain NA1000 CCNA GenBank locus ID. Data represent triplicate purifications of gsrN(37)⸬PP7hp and duplicate PP7hp⸬3’GsrN control purifications. Log adjusted p-values of zero are plotted as 10^-260^. (D) Sliding-window analysis of transcripts that co-purify with gsrN(37)⸬PP7hp (Figure 5-source data 2). Points represent 25-bp genome windows. RPKM values for each window were estimated by EDGE-pro; p-values were estimated by DESeq. Windows that map to genes identified in (C) are blue. Orange indicates windows with significant and highly abundant differences in mapped reads between gsrN(37)⸬PP7hp fractions and the PP7hp⸬gsrN-3’ negative control fractions. Dashed red lines denote cut-off value for windows enriched in the gsrN(37)⸬PP7hp fractions. Grey points within the dashed red lines are signal that mapped to rRNA. (E) Predicted loops in GsrN accessible for mRNA target base pairing are highlighted in yellow. A putative mRNA target site complementary to a cytosine-rich tract in the 5’ GsrN loop is represented as a sequence logo. Similar logo was generated for the sequences that mapped to the 2^nd^ exposed region of GsrN. Logo was generated from IntaRNA predicted GsrN-binding sites in transcripts enriched in the gsrN(37)⸬PP7hp pull-down. 5’ binding motif is present in 32 of the transcripts identified in (C) and (D) and 3’ binding motif is present in 27 of the transcripts identified in (C) and (D). (F) Density of reads that co-purified with gsrN(37)⸬PP7hp (blue) and PP7hp⸬gsrN-3’ (red) and mapped to *katG.* Read density in each dataset represents read coverage at each nucleotide divided by the number of million reads mapped in that data set. Data represent mean ± SD of replicate purifications.

The PP7hp aptamer inserted at *gsrN* nucleotide 37 (GsrN(37)⸬PP7hp) was selected as the bait to identify molecular partners that co-purify with GsrN. The pull-down fraction was compared to a negative control pull-down from cells expressing PP7hp fused to the last 50 nucleotides of GsrN including its intrinsic terminator (PP7hp⸬GsrN-3’) (Figure 5A). Northern blots demonstrated GsrN-PP7hp fusion transcripts were enriched in our purification (Figure 5B). Electrophoretic separation of the eluate followed by silver staining revealed no significant protein differences between GsrN(37)⸬PP7hp and the negative control (data not shown). We identified and quantified co-eluting RNAs by RNA-seq.

We employed two approaches to identify RNAs enriched in GsrN(37)⸬PP7hp fractions relative to the negative control fractions. A conventional RNA-seq pipeline (Tjaden, 2015) quantified mapped reads within annotated gene boundaries as a first pass (Figure 5C and Figure 5- source data 1). To capture reads in non-coding and unannotated regions, and to analyze reads unevenly distributed across genes, we also developed a sliding window analysis approach. Specifically, we organized the *Caulobacter* genome into 25 base-pair windows and quantified mapped reads in each window using the EDGE-pro/DESeq pipeline (Anders and Huber, 2010; Magoc et al., 2013). Together these two quantification strategies identified several mRNA, sRNAs, and untranslated regions enriched in the GsrN(37)⸬PP7hp pull-down fraction (Figure 5D and Figure 5- source data 2). We applied IntaRNA (Mann et al., 2017) to identify potential binding sites between GsrN and the enriched co-purifying RNAs. Of the 67 analyzed enriched genes and regions, 32 of the predicted RNA-RNA interactions involved the cytosine-rich 5’ loop in the predicted secondary structure of GsrN (Figure 5E and Figure 5- source data 3). A sequence logo (Crooks et al., 2004) of the predicted target mRNA binding sites is enriched with guanosines (Figure 5E), consistent with a model in which 6 tandem cytosines in the 5’ loop of GsrN determine target mRNA recognition. 27 of the predicted RNA-RNA interactions involved the 3’ exposed region of GsrN. The remaining 8 enriched genes and regions did not have a significant binding site prediction with GsrN.

Transcripts enriched in the GsrN(37)⸬PP7hp fraction encode proteins involved in proteolysis during envelope stress, enzymes required for envelope biogenesis, cofactor and nucleotide anabolic enzymes, and transport proteins (Table 1). *sigT* and its anti-σ factor, *nepR*, were also enriched in the GsrN(37)⸬PP7hp fraction, though we found no evidence for regulation of σ^T^/NepR by GsrN (Figure 1- figure supplement 2D). We observed significant enrichment of rRNA in the GsrN(37)⸬PP7hp fractions; the functional significance of this signal is not known (Figure 5D). *katG*, which encodes the sole catalase-peroxidase in the *Caulobacter* genome (Marks et al., 2010), was among the highly enriched mRNAs in our pull-down. Specifically, reads mapping to the first 60 nucleotides of *katG* including the 5’ leader sequence and the first several codons of the open reading frame were enriched in the GsrN(37)⸬PP7hp pull-down fraction relative to the negative control (Figure 5F). *katG* was an attractive GsrN target to interrogate the mechanism by which GsrN determines cell survival under hydrogen peroxide stress.

**Table 1.** RNAs that co-elute with GsrN-PP7hp.

| Gene Locus ID | Gene Name | log<sub>2</sub> Fold | Identification Method | Regions(s) | Description |
| --- | --- | --- | --- | --- | --- |
| CCNA_00167 | - | 4.56, 6.95 | Rockhopper, Sliding window | 179311-180120 (+), 179500-179550 (+, I, S) | metallophosphatase family protein |
| CCNA_00416 | - | 7.2 | Sliding window | 429625-429725 (-, I, S) | conserved hypothetical membrane protein |
| CCNA_00587 | - | 4.87 | Sliding window | 616250-616300 (+, I, S) | alpha/beta hydrolase family protein |
| CCNA_00882 | - | 4.61 | Sliding window | 962875-962925 (-, U, S) | hypothetical protein |
| CCNA_00894 | - | 4.29 | Sliding window | 974800-974850 (+, I, S) | 1-hydroxy-2-methyl-2-(E)-butenyl 4-diphosphate synthase |
| CCNA_00897 | - | 3.2 | Rockhopper | 976013-976177 (+) | hypothetical protein |
| CCNA_00913 | - | 7.64, 7.80 | Rockhopper, Sliding window | 993033-993209 (-), 993175-993225 (-, I, S) | hypothetical protein |
| CCNA_00930 | - | 3.72, 6.98, 3.81 | Rockhopper, Sliding window, Sliding window | 1006253-1006870 (+), 1006275-1006425 (+, I, S), 1006475-1006650 (+, I, S) | riboflavin synthase alpha chain |
| CCNA_01024 | - | 3.32 | Rockhopper | 1111617-1112111 (-) | hypothetical protein |
| CCNA_01058 | - | 5.81 | Sliding window | 1159075-1159125 (-, D, S) | helix-turn-helix transcriptional regulator |
| CCNA_01154 | - | 3.45, 6.81 | Rockhopper, Sliding window | 1257902-1258591 (+), 1257975-1258025 (+, I, S) | conserved hypothetical protein |
| CCNA_01303 | - | 5.87, 5.30, 8.22 | Rockhopper, Sliding window, Sliding window | 1430061-1430900 (+), 1430550-1430625 (+, I, S), 1430650-1430725 (+, I, S) | conserved hypothetical protein |
| CCNA_01304 | - | 2.9 | Rockhopper | 1431129-1431329 (+) | hypothetical protein |
| CCNA_01335 | - | 2.99 | Sliding window | 1448600-1448650 (-, I, S) | ABC-type multidrug transport system, ATPase component |
| CCNA_01344 | - | 4.62 | Sliding window | 1458550-1458725 (+, I, S) | conserved hypothetical protein |
| CCNA_01584 | - | 3.14 | Sliding window | 1699675-1699725 (+, I, A) | multimodular transpeptidase-transglycosylase PBP 1A |
| CCNA_01660 | - | 4.41, 6.21 | Rockhopper, Sliding window | 1781219-1781911 (-), 1781350-1781575 (-, I, S) | conserved hypothetical protein |
| CCNA_01966 | - | 11.3 | Sliding window | 2110225-2110275 (-, I, A) | vitamin B12-dependent ribonucleotide reductase |
| CCNA_01996 | - | 9.15, 8.86 | Rockhopper, Sliding window | 2142908-2143687 (-), 2143625-2143700 (-, I, S) | undecaprenyl pyrophosphate synthetase |
| CCNA_02034 | - | 7.24 | Sliding window | 2178500-2178550 (+, I, S) | luciferase-like monooxygenase |
| CCNA_02064 | lpxC | 3.6 | Sliding window | 2215450-2215550 (-, I, S) | UDP-3-O-(3-hydroxymyristoyl) N-acetylglucosamine deacetylase |
| CCNA_02089 | - | 8.52 | Rockhopper | 2237967-2238341 (-) | hypothetical protein |
| CCNA_02217 | - | 4.02 | Rockhopper | 2364081-2364383 (-) | hypothetical protein |
| CCNA_02286 | - | 3.26 | Sliding window | 2435450-2435500 (-, I, S) | hypothetical protein |
| CCNA_02595 | - | 6.85 | Sliding window | 2743525-2743625 (-, U, S) | Zn finger TFIIIB-family transcription factor |
| CCNA_02758 | - | 2.93 | Rockhopper | 2921763-2922152 (+) | hypothetical protein |
| CCNA_02761 | - | 3.65 | Rockhopper | 2923673-2923918 (+) | hypothetical protein |
| CCNA_02846 | - | 5.44, 8.60 | Sliding window, Sliding window | 3000100-3000175 (-, I, S), 2999225-2999275 (-, I, S) | DegP/HtrA-family serine protease |
| CCNA_02860 | - | 3.70, 4.78 | Rockhopper, Sliding window | 3012116-3013060 (-), 3012500-3012550 (-, I, S) | DnaJ-class molecular chaperone |
| CCNA_02975 | - | 6.34 | Sliding window | 3130300-3130375 (-, I, A) | excinuclease ABC subunit C |
| CCNA_02987 | - | 7.26 | Sliding window | 3142700-3142800 (-, I, A) | hypothetical protein |
| CCNA_02997 | cspA | 3.61 | Rockhopper | 3152607-3152816 (-) | cold shock protein CspA |
| CCNA_03002 | - | 6.03,<br>4.48 | Rockhopper,<br>Sliding window | 3155705-3156322 (-),<br>3155750-3155800 (-, I, S) | CDP-diacylglycerol-glycerol-3-phosphate 3<br>phosphatidyltransferase |
| CCNA_03105 | - | 9.15 | Sliding window | 3255775-3255850 (-, I, S) | DnaJ domain protein |
| CCNA_03113 | - | 3.50,<br>5.40 | Rockhopper,<br>Sliding window | 3263780-3264499 (-),<br>3264400-3264450 (-, I, S) | membrane-associated phospholipid<br>phosphatase |
| CCNA_03138 | katG | 3.35 | Sliding window | 3286000-3286050 (+, I, S) | peroxidase/catalase katG |
| CCNA_03176 | - | 2.83 | Rockhopper | 3335155-3335445 (-) | nucleotidyltransferase |
| CCNA_03338 | tolB | 5.27 | Sliding window | 3519425-3519475 (-, I, S) | TolB protein |
| CCNA_03409 | - | 4.46,<br>5.44 | Rockhopper,<br>Sliding window | 3576740-3577696 (-),<br>3577550-3577600 (-, I, S) | alpha/beta hydrolase family protein |
| CCNA_03506 | - | 3.27,<br>4.10 | Rockhopper,<br>Sliding window | 3664090-3664677 (+),<br>3664100-3664175 (+, I, S) | putative transcriptional regulator |
| CCNA_03589 | sigT | 3.58 | Rockhopper | 3743953-3744558 (-) | RNA polymerase EcfG family sigma<br>factor sigT |
| CCNA_03590 | nepR | 3.43,<br>3.50 | Rockhopper,<br>Sliding window | 3744561-3744746 (-),<br>3744675-3744725 (-, I, S) | anti-sigma factor NepR |
| CCNA_03590,<br>CCNA_03589 | nepR,<br>sigT | 4.42 | Sliding window | 3744500-3744575 (-, O, S) | anti-sigma factor NepR, RNA<br>polymerase EcfG family sigma factor<br>sigT |
| CCNA_03617 | - | 3.5 | Rockhopper | 3772262-3772717 (+) | Copper(I)-binding protein |
| CCNA_03618,<br>CCNA_03617 | -, - | 6.99 | Sliding window | 3772700-3772750 (+, O, S) | SCO1/SenC family protein, Copper(I)-<br>binding protein |
| CCNA_03681 | - | 5.11 | Sliding window | 3843700-3843750 (-, U, S) | ABC transporter ATP-binding protein |
| CCNA_03825 | - | 3.63 | Rockhopper | 3991412-3991774 (-) | hypothetical protein |
| CCNA_03825,<br>CCNA_03826 | -, - | 8.01 | Sliding window | 3991750-3991825 (-, O, S) | hypothetical protein, conserved<br>hypothetical protein |
| CCNA_03826 | - | 3.71 | Rockhopper | 3991771-3992325 (-) | conserved hypothetical protein |
| CCNA_03888 | - | 2.99 | Rockhopper | 761965-762324 (+) | conserved hypothetical protein |
| CCNA_03976 | - | 3.57 | Rockhopper | 2923462-2923683 (+) | hypothetical protein |
| CCNA_R0016 | - | 8.53 | Rockhopper | 844332-844401 (+) | small non-coding RNA |
| CCNA_R0035 | - | 6.64 | Rockhopper | 1549367-1549443 (+) | tRNA-Pro |
| CCNA_R0044 | - | 4.82 | Rockhopper | 2059848-2059942 (-) | complex medium expressed sRNA |
| CCNA_R0061 | - | 4.65 | Sliding window | 2800475-2800525 (-, I, S) | RNase P RNA |
| CCNA_R0089 | - | 3.3 | Sliding window | 3874375-3874425 (+, U, S) | tRNA-Ala |
| CCNA_R0100 | - | 4.4 | Rockhopper | 165492-165575 (+) | small non-coding RNA |
| CCNA_R0108 | - | 4.27 | Rockhopper | 472905-472973 (+) | small non-coding RNA |
| CCNA_R0180 | - | 5.14 | Rockhopper | 3266851-3266937 (-) | small non-coding RNA |

### GsrN base pairing to the 5’ leader of *katG* activates *katG* translation, and enhances peroxide stress survival

Most bacterial sRNAs regulate gene expression at the transcript and/or protein levels through Watson-Crick base pairing with the 5’end of their mRNA targets (Wagner and Romby, 2015). We sought to test whether GsrN affected the expression of *katG.* GsrN did not effect *katG* transcription in exponential or stationary phases, or in the presence of peroxide as measured by a *katG-lacZ* transcriptional fusion (Figure 6- figure supplements 1A-C). However, *katG* is transcriptionally regulated by the activator OxyR, which binds upstream of the predicted *katG* −35 site (Italiani et al., 2011). To decouple the effects of OxyR and GsrN on *katG* expression we generated a strict *katG* translational reporter that contains the mRNA leader of *katG* fused to *lacZ (katG-lacZ)* constitutively expressed from a σ^RpoD^-dependent promoter. In both exponential and stationary phases, *katG*-*lacZ* activity is reduced in Δ*gsrN* and enhanced in *gsrN^++^* strains compared to wild type (Figure 6- figure supplements 1D,F). Hydrogen peroxide exposure did not affect *katG*-*lacZ* activity (Figure 6- figure supplement 1E). We conclude that GsrN enhances KatG protein expression, but not *katG* transcription.

We then used this translational reporter to investigate a predicted binding interaction between the unpaired 5’ loop of GsrN and a G-rich region at the 5’ end of the *katG* transcript. Specifically, the first 7 nucleotides of *katG* mRNA (Zhou et al., 2015) is complementary to 7 nucleotides in the single-stranded 5’ loop of GsrN, including 4 of the 6 cytosines (Figure 6A). We disrupted this predicted base pairing, mutating 5 of the 7 nucleotides in the putative *katG* target site and GsrN interaction loop. These mutations preserved GC-content, but reversed and swapped (RS) the interacting nucleotides (Figure 6A). We predicted that pairs of wild-type and RS mutant transcripts would not interact, while base pairing interactions would be restored between RS mutant pairs.

**Figure 6.**
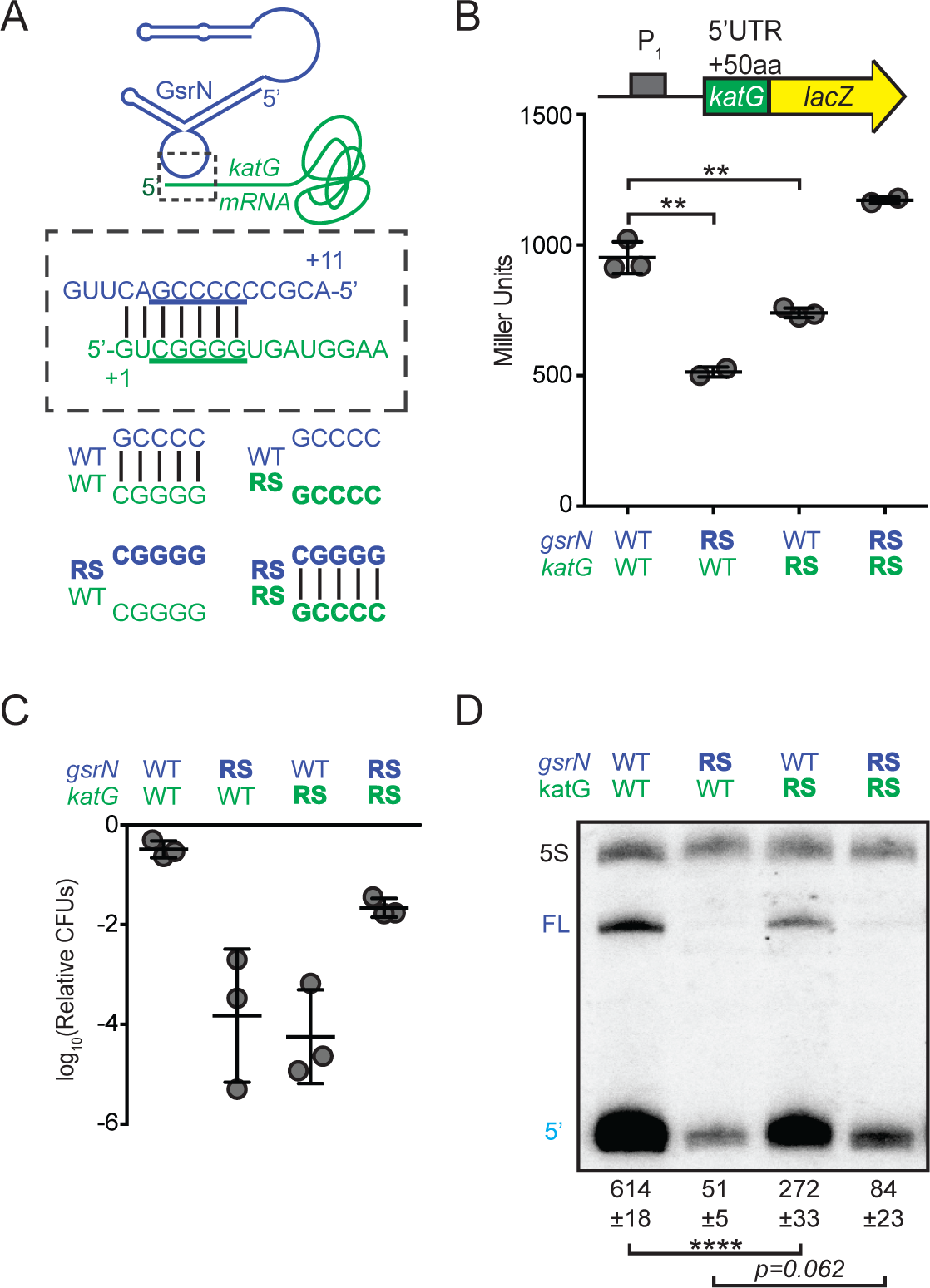
GsrN base pairs with the 5’ leader of *katG* mRNA and enhances KatG expression. (A) Predicted interaction between GsrN (blue) and *katG* mRNA (green), with base-pairing shown in dashed box. Wild-type (WT) and reverse-swapped (RS) mutation combinations of the underlined bases are outlined below. (B) Translation from *katG* and *katG-RS* reporters in Δ*gsrN* strains expressing 3*gsrN* (WT) or 3*gsrN(RS)* (RS). Measurements were taken from exponential phase cultures. Bars represent mean ± SD of independent cultures (points) and p-value estimated by Student’s t-test (C) Relative hydrogen peroxide survival of RS strains. Δ*gsrN* strains expressing 3*gsrN* or 3*gsrN(RS)* and encode *katG* or *katG(RS)* alleles. Bars represent mean ± SD from 3 independent experiments (points). (D) Northern blot of total RNA from strains in (C) collected in exponential phase hybridized with probes complementary to 5’ end of GsrN and 5S rRNA. Quantification is mean ± SD normalized signal from 3 independent experiments and p-value estimated by Student’s t-test.

Mutating the predicted target site in the *katG* 5’ leader ablated GsrN-dependent regulation of the *katG*-*lacZ* translational reporter (Figure 6- figure supplement 2A); expression was reduced to a level similar to Δ*gsrN*. We further tested this interaction by assessing the effect of the reverse-swapped *gsrN(RS)* allele on the expression of *katG-lacZ.* However, GsrN(RS) was unstable; total GsrN(RS) levels were ≈10-fold lower than wild-type GsrN (Figure 6- figure supplements 3A,B). To overcome GsrN(RS) instability, we inserted a plasmid with three tandem copies of gsrN(RS), 3gsrN(RS), into the *vanA* locus in a Δ*gsrN* background, which increased steady-state levels of GsrN(RS) approximately 4-fold (Figure 6- figure supplements 3A,B). *katG* target site or GsrN recognition loop mutations significantly reduced *katG*-*lacZ* expression (Student’s t-test, *p*=*0.0026* and *p*=*0.0046*, respectively). Compensatory RS mutations that restored base pairing between the *katG* target site and the GsrN loop rescued *katG*-*lacZ* expression (Figure 6B).

To assess the physiological consequence of mutating the G-tract in the *katG* mRNA leader and the GsrN C-rich loop, we replaced wild-type *katG* on the chromosome with the *katG(RS)* allele in both the Δ*gsrN*+3*gsrN* and Δ*gsrN*+3*gsrN*(RS) backgrounds, and measured survival after hydrogen peroxide exposure. Both *katG(RS)* and *gsrN(RS)* mutants had survival defects (Figure 6C and Figure 6- figure supplement 3C). Strains harboring the *katG(RS)* allele phenocopy the survival phenotype of Δ*gsrN* under peroxide stress. While *katG(RS)* survival is compromised, the defect is not as large as a strain missing *katG* completely (Δ*katG*) (Figure 6C and Figure 6- figure supplement 2C). Expressing *gsrN(RS)* in one, three, or six tandem copies did not complement the peroxide survival defect of Δ*gsrN* (Figure 6- figure supplement 3C). A strain expressing *katG(RS)* and *gsrN(RS)*, which restores base pairing between the GsrN 5’ loop and the *katG* 5’ leader, rescued hydrogen peroxide stress survival (Figure 6C). The protective effect of overexpressing *gsrN* is lost when *katG* is deleted, and overexpression of *katG* rescues the survival defect of Δ*gsrN* after peroxide treatment. We conclude that *katG* is necessary and sufficient to protect the cell from hydrogen peroxide (Figure 6- figure supplement 2C).

Given differences in steady state levels of GsrN and GsrN(RS), we postulated that the capacity of GsrN to interact with its targets influences its stability *in vivo*. Indeed, mutation of the *katG* target site reduced GsrN by more than 2-fold (Student’s t-test, *p*<*0.0001*). The compensatory *katG(RS)* allele partially restored stability to GsrN(RS) (Figure 6D). *katG(RS)* mutation or *katG* deletion did not influence *gsrN* transcription (Figure 6- figure supplement 2B). Thus, we attribute the differences in steady-state levels of the GsrN alleles to their ability to interact with mRNA targets via the 5’ C-rich loop.

### GsrN enhances KatG expression and stabilizes *katG* mRNA *in vivo* in the presence of peroxide

To assess the relative effects of GsrN on *katG* transcript and protein levels *in vivo*, we directly measured both by dot blot and Western blot, respectively. In untreated and peroxide treated cultures, *katG* transcript levels trended lower in Δ*gsrN* and higher in *gsrN^++^* compared to wild type. These differences are not statistically significant (Student’s t-test, *p*=*0.39)* in untreated cultures; however, KatG protein tagged with the M2 epitope was reduced 2-fold in Δ*gsrN* lysates relative to wild-type in untreated cultures (Student’s t-test, *p*<*0.0001*) (Figure 7B). Since GsrN does not influence *katG* transcription in untreated cultures (Figure 6- figure supplements 1A,C), GsrN may enhance KatG translation *in vivo.*

**Figure 7.**
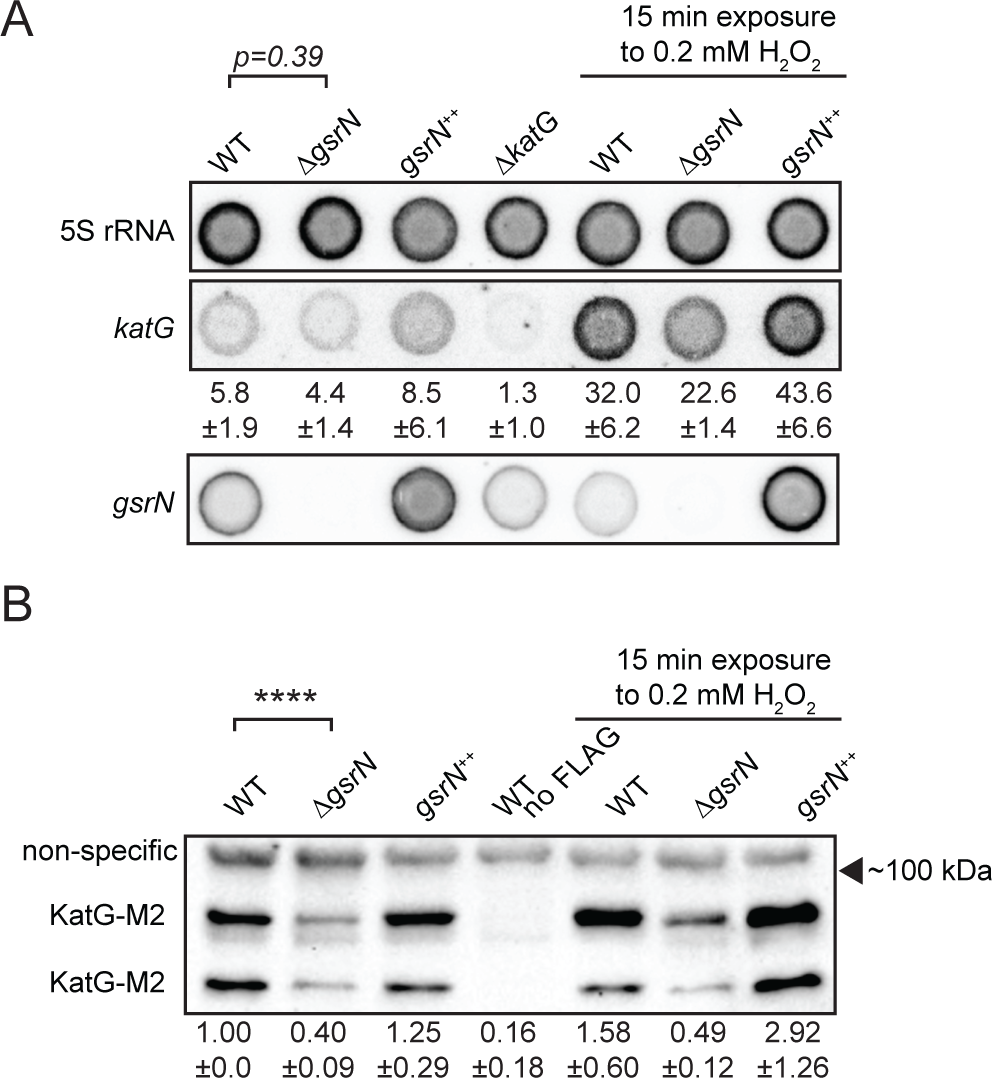
GsrN affects KatG and *katG* mRNA levels *in vivo.* (A) Dot blot of total RNA of *gsrN* and *katG* mutants grown to early stationary phase (OD_660_ 0.85-0.9). Samples on right were treated with 0.2 mM hydrogen peroxide before RNA extraction. Blots were hybridized with *katG* mRNA, GsrN or 5S rRNA probes. *katG* mRNA signal normalized to 5S rRNA signal is quantified (mean ± SD, n=3, p-value estimated with Student’s t-test). (B) Immunoblot of KatG-M2 fusion in wild type, Δ*gsrN*, and Δ*gsrN^++^* strains in the presence and absence of peroxide stress probed with α-FLAG antibody. KatG migrates as two bands as previously reported (Italiani et al., 2011). Normalized KatG-M2 signal (mean ± SD, n=4, **** p<0.0001 Student’s t-test) is presented below each lane. Arrow indicates position of 100 kDa molecular weight marker.

Steady-state *katG* transcript levels differ significantly between Δ*gsrN* and *gsrN^++^* in peroxide treated cultures (Student’s t-test, *p*<*0.01*) (Figure 7A). KatG protein tagged with the M2 epitope was reduced 3-fold in peroxide treated cells in Δ*gsrN* lysates relative to wild-type; KatG-M2 levels in *gsrN^++^* were increased in both untreated and peroxide treated cells (Figure 7B). These data support a model whereby GsrN enhances KatG protein expression in the presence of peroxide by stabilizing *katG* mRNA and/or promoting *katG* translation.

### GsrN is a general regulator of stress adaptation

In the GsrN⸬PP7hp pull-down fraction, we observed enrichment of multiple RNAs in addition to *katG* (Figure 5C,D). This suggested that GsrN may have regulatory roles beyond mitigation of peroxide stress. To globally define genes that are directly or indirectly regulated by GsrN, we performed RNA-seq and LC-MS/MS measurements on wild-type, Δ*gsrN* and *gsrN^++^* strains (Figure 8A and Figure 8- source data 1). We identified 40 transcripts, including *gsrN*, with significant differences in mapped reads between the Δ*gsrN* and *gsrN^++^* samples (Figure 8- figure supplement 1A and Figure 8- source data 2). 11 proteins had significant label free quantitation (LFQ) differences (FDR<0.05) between *gsrN^++^* and Δ*gsrN* (Figure 8- figure supplement 1B and Figure 8- source data 3). Most genes identified as significantly regulated by transcriptomic and proteomic approaches did not overlap. Nonetheless, these data provide evidence that GsrN can function as both a positive and negative regulator of gene expression, either directly or indirectly.

**Figure 8.**
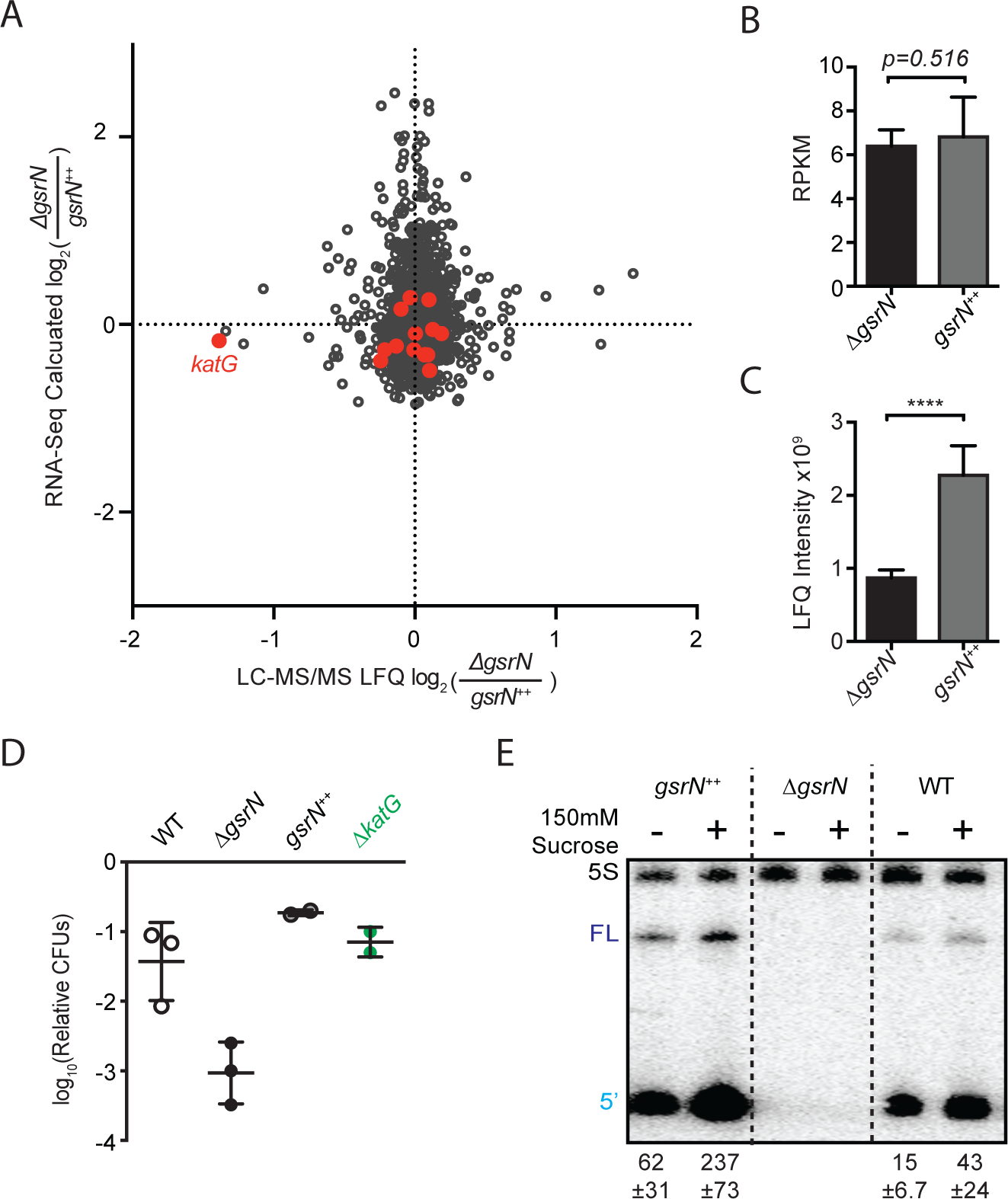
GsrN is a global regulator of stress physiology. (A) Transcriptomic and proteomic analysis of Δ*gsrN* (deletion) and *gsrN^++^* (overexpression) cultures in early stationary phase (Figure 8- source data 1). Only genes detected in both analyses are plotted. Red indicates transcripts that co-purify with GsrN-PP7hp (Figures 5C,D). (B) *katG* transcript from Δ*gsrN* and *gsrN^++^* cells quantified as reads per kilobase per million mapped (RPKM). Data represent mean ± SD of 5 independent samples. Significance was evaluated with the Wald test. (C) Label free quantification (LFQ) intensities of KatG peptides from Δ*gsrN* and *gsrN^++^* cells (mean ± SD, n=3; **** p<0.0001 Student’s t-test). (D) Hyperosmotic stress survival of wild type, Δ**gsrN*, gsrN*^++^, and Δ*katG* cells relative to untreated cells. Stress was a 5 hour treatment with 300 mM sucrose. Data represent mean ± SD from 2 independent experiments (points). (E) Northern blot of total RNA from wild type, Δ*gsrN*, and *gsrN*^++^ cultures with or without 150 mM sucrose stress. Blots were hybridized with GsrN and 5S rRNA probes. Normalized mean ± SD of total GsrN signal from 3 independent samples is quantified.

Importantly, RNA-seq and proteomics experiments validated *katG* as a regulatory target of GsrN. *katG* transcript levels measured by RNA-seq were not significantly different between Δ*gsrN* and *gsrN^++^* strains (Figure 8B), consistent with our dot blot measurements of unstressed cultures (Figure 7A). Conversely, steady-state KatG protein levels estimated from our LC-MS/MS experiments were significantly reduced in Δ*gsrN*, consistent with our Western analysis of KatG protein (Figures 8C,5F). *katG* was the only gene that that was significantly enriched in the pull-down and differentially expressed in the proteomic studies (Figure 8A). These results provide additional evidence that *katG* is a major target of GsrN, and that GsrN functions to enhance KatG expression at the post-transcriptional level.

Given our transcriptomic and proteomic datasets, we reasoned that GsrN may contribute to other phenotypes associated with deletion of the GSR sigma factor, *sigT*. Indeed, the Δ*gsrN* mutant has a survival defect after exposure to hyperosmotic stress, similar to Δ*sigT* (Figure 8D). As we observed for peroxide stress, overexpression of *gsrN* protects cells under this physicochemically-distinct condition. Hyperosmotic stress survival does not require *katG* (Figure 8D), providing evidence that a separate GsrN regulatory target mediates this response. Unlike hydrogen peroxide (Figure 7A), hyperosmotic stress induces GsrN expression (Figure 8E). This is consistent with previous transcriptomic studies in *Caulobacter* in which hyperosmotic stress, but not peroxide stress, activated GSR transcription (Alvarez-Martinez et al., 2007). GsrN transcription is also significantly enhanced in stationary phase cultures relative to logarithmic phase cultures (Figure 1- figure supplement 2E). Though its functional role under this condition remains undefined, it has been reported that *katG* is a genetic determinant of stationary phase survival (Steinman et al., 1997).

### σ^EcfG^-regulated sRNAs are prevalent across the Alphaproteobacterial clade

The GSR system is broadly conserved in Alphaproteobacteria. Given the importance of GsrN as a post-transcriptional regulator of the *Caulobacter* GSR, we reasoned that functionally-related sRNAs might be a conserved feature of the GSR in this clade. To identify potential orthologs of *gsrN*, we surveyed the genomes of Alphaproteobacteria that encoded regulatory components of the GSR system and for which transcriptomic data were publically available.

We initially searched for GsrN-related sequences using BLASTn (Altschul et al., 1990). Hits to GsrN were limited to the Caulobacteraceae family, including the genera *Caulobacter, Brevundimonas*, and *Phenylobacterium.* The 5’ C-rich loop of homologs identified in this family had the highest level of conservation compared to other regions of secondary structure (Figure 9B). Predicted *gsrN* homologs are often proximal to the genes encoding the core GSR regulators (*ecfG*/*sigT, nepR* and *phyR*) (Figure 9A). *C. crescentus* is a notable exception where *gsrN* is positioned distal to the GSR locus. Therefore, we used genome position as a key parameter to identify additional GsrN or GsrN-like RNAs in Alphaproteobacteria outside of Caulobacteraceae.

Our search for GsrN focused on three parameters: evidence of intergenic transcription, identification of a near-consensus σ^EcfG^ site in the promoter region, and proximity to the *sigT*-*phyR* chromosomal locus. Based on our criteria, we identified a set of putative GsrN homologs in the Rhizobiaceae family (Jans et al., 2013; Kim et al., 2014; Valverde et al., 2008) (Figure 9A). The predicted secondary structure of these putative GsrN homologues has features similar to GsrN from Caulobacteraceae. Specifically, there is an exposed cytosine-rich loop at the 5’ end (Figure 9C).

**Figure 9.**
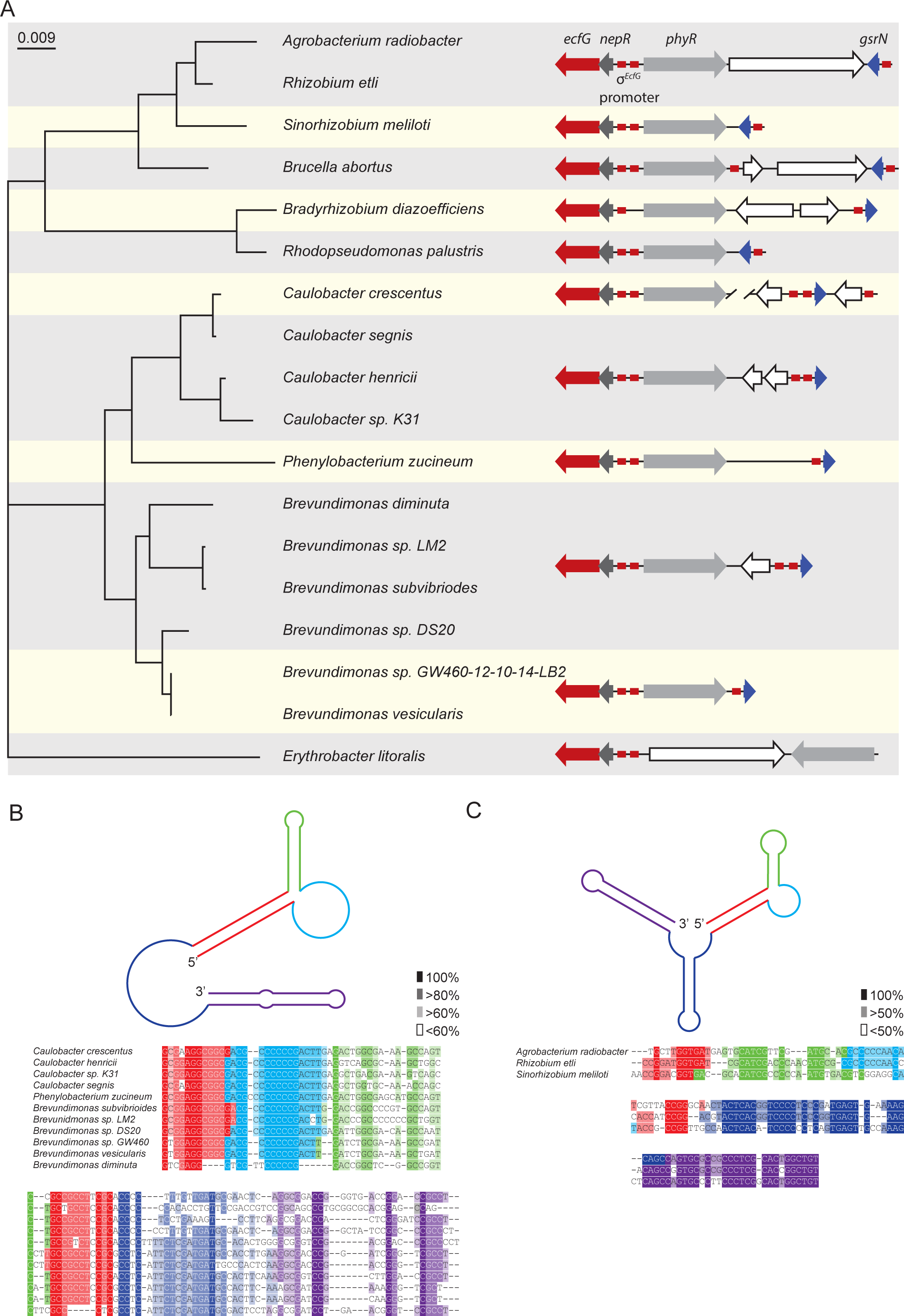
Conserved features of GsrN homologues. (A) Locus diagrams showing predicted *gsrN* homologs in several Alphaproteobacteria. Tree was constructed from the 16s rRNA sequences of each strain where *Erythrobacter litoralis* (for which there is no apparent gsrN-like gene) was the out-group. Red arrows represent *ecfG*, dark gray arrows represent *nepR*, red boxes represent the conserved σ^ecfG^-binding site, light gray arrows represent *phyR*, and dark blue arrows represent *gsrN* (or its putative homologs). The prediction of GsrN orthologs in the Caulobacteraceae (*Caulobacter, Brevundimonas*, and *Phenylobacterium*) was based on a BLASTn search (Altschul et al., 1990). The prediction of GsrN in *Rhizobium etli, Sinorhizobium meliloti*, and *Brucella abortus* was based on evidence of expression in published transcriptome data, proximity to the GSR locus, and identification of a σ^ecfG^-binding site upstream of the gene. The prediction of *Agrobacterium radiobacter* was based on a BLASTn search of using the predicted GsrN sequence from *R. etli* as the query (Altschul et al., 1990). The prediction of *Rhodopseudomonas palustris* and *Bradyrhizobium diazoefficiens* is completely based on the proximity to the GSR locus and the presence of an upstream σ^ecfG^-binding site. (B) Diagram of predicted secondary structure of GsrN in other Caulobacteraceae is colored by secondary structure element. Colors highlighted in the sequence alignment correspond to the predicted secondary structure regions in the cartoon. Density of shading corresponds to conservation at that position. (C) Diagram of predicted secondary structure of predicted GsrN homologs in select Rhizobiaceae where the 5’ portion contains an unpaired 5’ G-rich loop (cyan) flanked by a small hairpin (green) and a stem loop involving the 5’ terminus (red).

**Figure 10.**
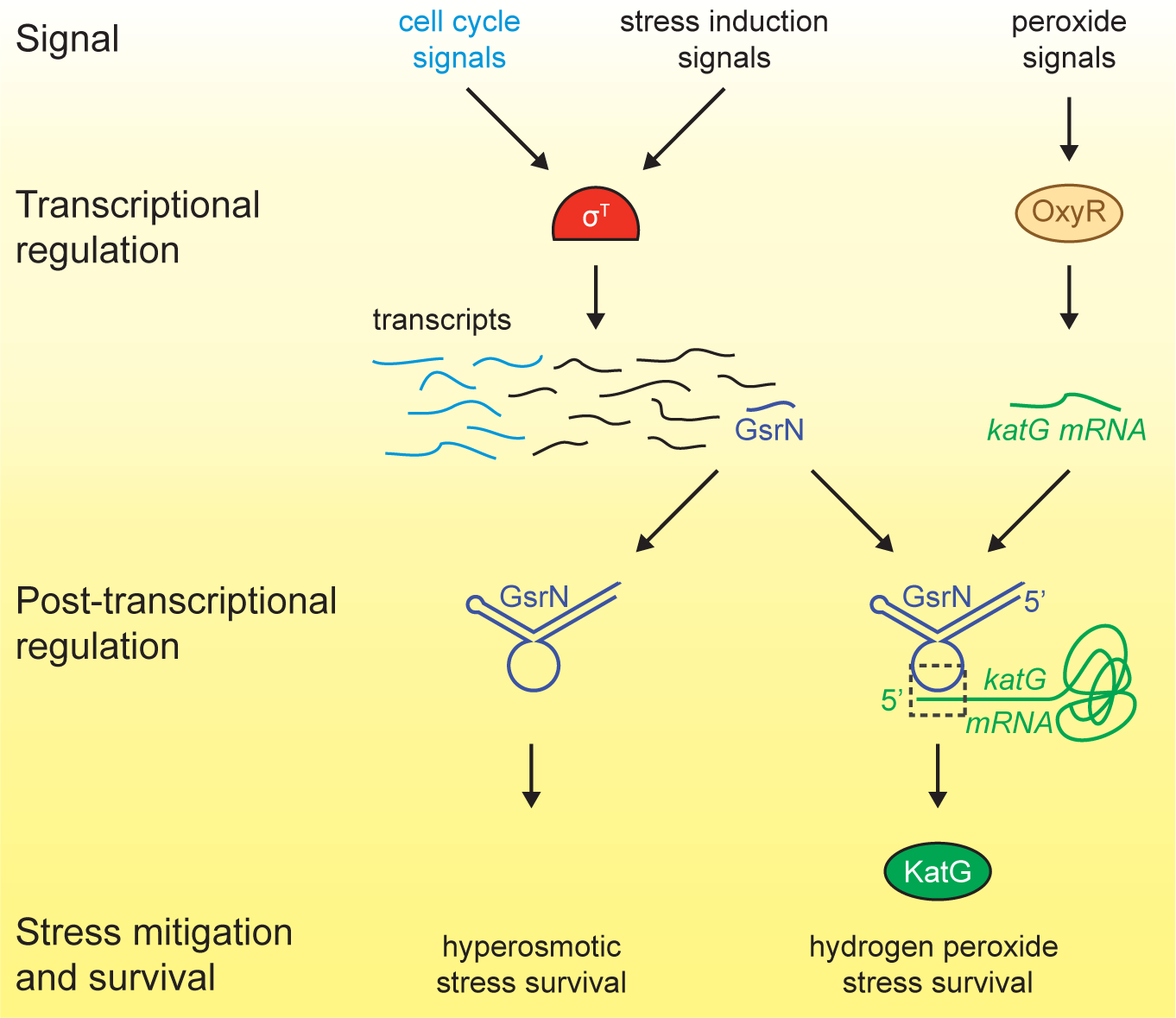
Regulatory architecture of the *Caulobacter* stress response systems. Expression of the GSR EcfG-sigma factor, *sigT* (σ^T^), and select genes in the GSR regulon is regulated as a function cell cycle phase. σ^T^-dependent transcription can be induced by certain signals (e.g. hyperosmotic stress), but is unaffected by hydrogen peroxide. Transcription of the sRNA, GsrN, is activated by σ^T^, and the cell cycle expression profile of *gsrN* is highly correlated with *sigT* and its upstream regulators. Transcription of the catalase/peroxidase *katG* is independent of σ^T^. GsrN dependent activation of KatG protein expression is sufficient to rescue the peroxide survival defect of a Δ*sigT* null strain. GsrN convenes a post-transcriptional layer of gene regulation that confers resistance to peroxide and hyperosmotic stresses.

## DISCUSSION

We sought to understand how GSR transcription determines cell survival across a spectrum of chemical and physical conditions. To this end, we developed a directed gene network analysis approach to predict genes with significant functional roles in the *Caulobacter* GSR. Our approach led to the discovery of *gsrN*, a small RNA of previously unknown function that is a major post-transcriptional regulator of stress physiology.

### Role of GsrN in mitigating hydrogen peroxide stress

Hydrogen peroxide can arise naturally from the environment and is also produced as an aerobic metabolic byproduct (Imlay, 2013). Our data provide evidence that σ^T^-dependent transcription of GsrN basally protects cells from hydrogen peroxide by enhancing KatG expression. Unlike the transcription factor OxyR, which induces *katG* expression in response to peroxide (Italiani et al., 2011), GsrN is not induced by peroxide treatment. KatG levels change by only a factor of two when *gsrN* is deleted or overexpressed, but we observe dramatic peroxide susceptibility and protection phenotypes as a function of *gsrN* deletion and overexpression, respectively. The survival phenotypes associated with subtle fold changes in KatG expression suggest that the capacity of KatG to detoxify endogenous sources of H_2_O_2_ is at or near its limit under normal cultivation conditions, similar to what has been postulated for *E. coli* (Imlay, 2013).

Expression of the ferritin-like protein, Dps, is controlled by σ^T^ and is reported to aid in the survival of *Caulobacter* under peroxide stress (de Castro Ferreira et al., 2016). The protective effect of Dps is apparently minimal under our conditions given that *a)* the peroxide survival defect of Δ*sigT* is rescued by simply restoring *gsrN* transcription (Figure 2B,C), and *b)* survival after peroxide exposure is determined almost entirely by modifying base-pairing interactions between GsrN and *katG* mRNA. This stated, the difference in hydrogen peroxide susceptibility between Δ*sigT* and Δ*gsrN* (Figure 1E) may be explained in part by the fact that *dps* is still expressed in Δ*gsrN*.

### Post-transcriptional gene regulation by GsrN is a central feature of the general stress response

Alternative sigma factor regulation is a major mechanism underlying transcriptional reprogramming in response to stress (Paget, 2015; Staron et al., 2009). Roles for sRNAs in regulation of environmental adaptation are also well-described (Storz et al., 2011). sRNAs have been implicated in regulation of the σ^S^-dependent general stress pathway of Enterobacteria (Mika and Hengge, 2014), which is functionally analogous to the Alphaproteobacterial GSR. sRNAs that function downstream of the σ^B^ general stress factor in Firmicutes have also been reported (Mader et al., 2016; Mellin and Cossart, 2012). In the case of the σ^S^ pathway of *E. coli*, different sRNAs function under different conditions to modulate *rpoS* regulatory output (Repoila et al., 2003). Our data define *Caulobacter* GsrN as a central regulator of stress physiology that has a major protective effect across distinct conditions. GsrN does not mitigate stress by feeding back to affect GSR dependent transcription. The effects we report here are, apparently, purely post-transcriptional and downstream of σ^T^. GsrN protects *Caulobacter* from death under hyperosmotic and peroxide stress conditions via genetically distinct post-transcriptional mechanisms (Figures 1,8). Thus, transcriptional activation of GsrN by σ^T^ initiates a downstream post-transcriptional program that directly affects multiple genes required for stress mitigation.

Quantitative proteomic studies (Figure 8A) demonstrate that GsrN activates and represses protein expression, either directly or indirectly. In the case of KatG, we have shown that GsrN is among the rare class of sRNAs (Frohlich and Vogel, 2009) that directly enhance protein expression (Figure 7B).

Our global and directed measurements of mRNA show that *katG* mRNA levels do not change significantly between Δ*gsrN* and *gsrN^++^* strains (Figure 7,8). However, in the presence of peroxide, we observe significant changes in *katG* mRNA that correlate with changes in KatG protein levels. Our data suggest a role for GsrN as a regulator of mRNA translation and, perhaps, mRNA stability. In this way, GsrN may be similar to the sRNAs, DsrA and RhyB (Lease and Belfort, 2000; Prevost et al., 2007). However, DsrA and RhyB function by uncovering ribosome-binding sites (RBS) in the leaders of their respective mRNA target. We are unable to predict a location for the RBS in the *katG* mRNA leader, but note that *katG* is among the 75% of open reading frames (ORFs) in *Caulobacter* that do not contain a canonical RBS (Schrader et al., 2014).

To our knowledge, no other sRNA deletion mutant has as dramatic a set of stress survival phenotypes as Δ*gsrN*. The target of GsrN that confers hyperosmotic stress protection remains undefined, but this phenotype is also likely regulated at the post-transcriptional level (Figure 8D). While the reported physiological effects of sRNAs are often subtle, GsrN provides a remarkable example of a single post-transcriptional regulator that exerts a strong influence on multiple, distinct pathways controlling cellular stress survival.

### On GsrN stability and processing

The roles of sRNAs in stress adaptation have been investigated in many species, and a number of molecular mechanisms underlying sRNA-dependent gene regulation have been described. We have uncovered a connection between mRNA target site recognition and GsrN stability that presents challenges in the characterization of GsrN regulatory mechanisms. Specifically, mutations in the *katG* mRNA leader affect steady-state levels of GsrN (Figure 6D). Given this result, one could envision scenarios in which changes in transcription of *katG* or some other direct GsrN target could broadly affect stress susceptibility by altering levels of GsrN and, in turn, the stability of other target mRNAs in the cell. In short, the concentrations of mRNA targets could affect each other via GsrN. Such effects should be considered when assessing mRNA target site mutations in this system and others.

GsrN is among a handful of sRNAs that are reported to be post-transcriptionally processed (Figure 3) (Papenfort and Vanderpool, 2015). Select PP7hp insertions resulted in reduced 5’ isoform formation; PP7hp insertion mutants with low 5’ isoform levels did not complement the peroxide viability defect of Δ*gsrN*. Processing to a short 5’ isoform may be necessary for GsrN to bind *katG* mRNA and regulate KatG expression. Alternatively, cleavage may not be required for function, and lack of complementation by certain hairpin insertion mutants may be due to PP7hp interfering with target recognition or simply reducing total levels of GsrN. Regardless, our data clearly show that GsrN is cleaved in a regular fashion to yield a 5’ isoform that is very stable in the cell (Figure 3- figure supplement 1) and is sufficient to protect *Caulobacter* from hydrogen peroxide treatment (Figure 4B). Our understanding of the role of RNA metabolism in sRNA-dependent gene regulation is limited, and GsrN provides a good model to investigate mechanisms by which mRNA target levels and sRNA and mRNA processing control of gene expression.

### *Caulobacter* GSR and the cell cycle

The transcription of *sigT*, *gsrN*, and several other genes in the GSR regulon are cell cycle regulated (Fang et al., 2013; Laub et al., 2000; McGrath et al., 2007; Zhou et al., 2015), with highest expression during the swarmer-to-stalked cell transition, when cells initiate DNA replication and growth (Figure 1C). GSR activation during this period potentially protects cells from endogenous stressors that arise from upregulation of anabolic systems required for growth and replication. In the future, it is of interest to explore the hypothesis that the GSR system provides both basal protection against endogenous stressors generated as a function of normal metabolism, and induced protection against particular stressors (e.g. hyperosmotic stress) encountered in the external environment.

## MATERIALS AND METHODS

### Experimental Model And Subject Details

#### Growth Media and Conditions

*C. crescentus* was cultivated on peptone-yeast extract (PYE)-agar (0.2% peptone, 0.1% yeast extract, 1.5% agar, 1 mM MgSO_4_, 0.5 mM CaCl_2_) (Ely, 1991) at 30°C. Antibiotics were used at the following concentrations on this solid medium: kanamycin 25 μg/ml; tetracycline 1 μg/ml; and chloramphenicol 2 μg/ml.

For liquid culture, *C. crescentus* was cultivated in either PYE or in M2X defined medium (Ely, 1991). PYE liquid: 0.2%(w/v) peptone, 0.1%(w/v) yeast extract, 1 mM MgSO_4_, and 0.5 mM CaCl_2_ autoclaved before use. M2X defined medium: 0.15% (w/v) xylose, 0.5 mM CaCl_2_, 0.5 mM MgSO_4_, 0.01 mM Fe Chelate, and 1x M2 salts, filtered with a 0.22 micron bottle top filter. One liter of 20x M2 stock was prepared by mixing 17.4g Na_2_HPO_4_, 10.6 KH_2_PO_4_, and 10g NH_4_Cl. To induce gene expression from the *vanA* promoter, 500 μM vanillate (final concentration) was added. Antibiotics were used at the following concentrations in liquid medium: kanamycin 5 μg/ml, tetracycline 1 μg/ml, nalidixic acid 20 μg/ml, and chloramphenicol 2 μg/ml.

For cultivation of *E. coli* in liquid medium, we used lysogeny broth (LB). Antibiotics were used at the following concentrations: ampicillin 100 μg/ml, kanamycin 50 μg/ml, tetracycline 12 μg/ml, and chloramphenicol 20 μg/ml.

#### Strain construction

All *C. crescentus* experiments were conducted using strain CB15 (Poindexter, 1964) and derivatives thereof. Plasmids were conjugated into CB15 (Ely, 1991) using the *E. coli* helper strain FC3 (Finan et al., 1986). Conjugations were performed by mixing the donor *E. coli* strain, FC3, and the CB15 recipient strain in a 1:1:5 ratio. Mixed cells were pelleted for 2 minutes at 15,000xg, resuspended in 100 μL, and spotted on a nonselective PYE-agar plate for 12-24 hours. Exconjugants containing the desired plasmid were spread on PYE agar containing the plasmid-specified antibiotic for selection. The antibiotic nalidixic acid (20 μg/ml) was used to counter-select against both *E. coli* strains (helper and plasmid donor).

Gene deletion and nucleotide substitution strains were generated using the integrating plasmid pNPTS138 (Ried and Collmer, 1987). pNPTS138 transformation and integration occurs at a chromosomal site homologous to the insertion sequence in pNPTS138. Exconjugants with pNPTS138 plasmids were selected on PYE agar plates with 5 μg/ml kanamycin; 20 μg/ml nalidixic acid selected against the *E. coli* donor strain. Single colony exconjugants were inoculated into liquid PYE or M2X for 6-16 hours in a rolling 30°C incubator for non-selective growth. Nonselective liquid growth allows for a second recombination event to occur, which either restores the native locus or replaces the native locus with the insertion sequence that was engineered into pNPTS138. Counter-selection for the second recombination of pNPTS138 was carried out on PYE agar with 3% (w/v) sucrose. This selects for loss of the *sacB* gene during the second crossover event. Colonies were subjected to PCR genotyping and/or sequencing to identify to confirm the allele replacement.

Other strains utilized in this study originate from (Herrou et al., 2010), (Purcell et al., 2007), and (Foreman et al., 2012).

The Δ*gsrN* strains and Δ*sigT*strains were complemented by introducing the gene at an ectopic locus (either *vanA* or *xylX*) utilizing the integrating plasmids: pMT552, pMT674, and pMT680. pMT674 and pMT680 carry a chloramphenicol resistance marker gene (*cat*) and pMT552 carries a kanamycin resistance marker gene (*npt1*) (Thanbichler et al., 2007). pMT552 and pMT674 integrate into the *vanA* gene and pMT680 integrates into the *xylX* gene. Transformation of ectopic complementation plasmids conjugated (as described earlier). Introduction of *gsrN* complementation was done in the reverse direction of the inducible promoters. Introduction of *katG* was done in-frame in the same direction of the inducible promoters.

Replicating plasmids pPR9TT and pRKlac290 were conjugated as previously described earlier. pPR9TT and pRKlac290 were selected using tetracycline and chloramphenicol, respectively.

pMal-MBP-PP7CPHis was transformed into *E. coli* Rosetta by electroporation and plated on LB plates with ampicillin 100 μg/ml.

#### Plasmid construction

Plasmid pNPTS138 was used to perform allele replacements and to generate gene deletions (Ried and Collmer, 1987; West et al., 2002). Primers for in-frame deletions and GeneBlocks (Gblocks) are listed in Supplementary File 1. Gene fragments were created by splice-overlap-extension and ligated into the digested pNPTS138 vector at restriction enzyme sites (HindIII, SpeI) or gene fragments were stitched together using Gibson assembly. pNPTS138 contains a *kan^R^* (*npt1*) antibiotic resistance marker and the counter-selectable marker gene *sacB*, which encodes levansucrase Plasmids for *gsrN* genetic complementation experiments carried wild-type or mutant *gsrN* alleles cloned antisense into a vanillate inducible(*vanA*)-promoter. An in-frame stop codon was designed at a restriction enzyme site downstream of the *vanA* promoter to ensure translational read-through of the *vanA* transcript did not disrupt *gsrN* transcription. Tandem *gsrN* alleles (overexpression by multiple copies of *gsrN*) were constructed using Gblocks with unique ends for Gibson assembly into pMT552. Plasmids for genetic complementation of the *katG* mutant were constructed by cloning *katG* in-frame with the vanillate and xylose-inducible promoters of pMT674 and pMT680, respectively, at the NdeI and Kpnl restriction sites. *katG* complementation plasmids did not include the 5’ untranslated region (UTR) of *katG.*

Beta-galactosidase transcriptional and translational reporters utilized pRKlac290 (Ely, 1991) and pPR9TT (Santos et al., 2001) replicating plasmids, respectively. Transcriptional reporters of *gsrN* contained upstream and promoter sequences of *gsrN* cloned into the EcoRI and HindIII sites of pPRKlac290. Translational reporters of *katG* contained the 191 nucleotides 3’ of the annotated *katG* transcriptional start site (Zhou et al., 2015) cloned into pPR9TT at HindIII and Kpnl.

Protein expression plasmid pMal was used to express a maltose binding protein (MBP) fused to the N-terminus of a Pseudomonas Phage 7 coat protein fused to a His-tag at its C-terminus (to generate MBP-PP7CP-His). The PP7CPHis protein sequence was amplified out of pET283xFlagPP7CPHis and inserted into pMal at SalI and EcoRI restriction sites. pET283xFlagPP7CPHis was a gift from Alex Ruthenburg and originates from Kathleen Collins (Addgene plasmid # 28174).

### Experimental Method Details

#### Hydrogen peroxide/osmotic stress assays

Liquid cultures were passaged several times before stress treatment to insure that population growth rate and density was as consistent as possible prior to addition of hydrogen peroxide (oxidative stress) or sucrose (hyperosmotic stress). Briefly, starter cultures were inoculated in liquid M2X medium from colonies grown on PYE-agar plates. Cultures were grown overnight at 30°C in a rolling incubator. Overnight cultures were then diluted back to an optical density reading of 0.05 at 660 nm (OD_660_=0.05) and grown in a rolling incubator at 30°C for 7-10 hours. After this period, cultures were re-diluted with M2X to OD_660_=0.025 and grown for 16 hours at 30°C in a rolling incubator. After this period, OD_660_ was consistently 0.85-0.90. These cultures were then diluted to OD_660_=0.05 and grown for 1 hour and split into two tubes. One tube received stress treatment and the other tube was untreated. Treated cultures were subjected to either hydrogen peroxide or sucrose.

For stress treatment, we used a freshly prepared 10 mM H_2_O_2_ solution diluted from a 30% (w/w) stock bottle (stock never more than 3 months old) or a stock of 80% (w/v) sucrose. The amount of 10 mM H_2_O_2_ added for stress perturbation depended on the volume of the culture and the desired final concentration of H_2_O_2_. Final volumes assessed in our studies are described for each experiment throughout this manuscript.

Treated cultures and untreated cultures were subsequently tittered (10 μL sample in 90 μL of PYE) by initially diluting into 96-well plates. 5 μL spots from each dilution were plated on PYE-agar. Once spots dried, plates were incubated at 30°C for 2 days. Clearly visible colonies begin to form after 36 hours in the incubator.

The difference in colony forming units (CFU) between treated and untreated cultures was calculated using the following formula:

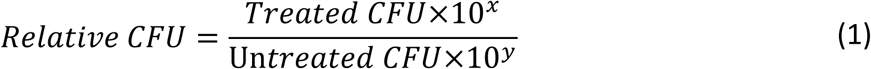

Where *x* represents the countable (resolvable) dilution in which colonies are found in the treated sample dilution series and y represents the untreated sample dilution.

#### β-galactosidase gene expression reporter assays

To assess reporter gene expression, liquid cultures were passaged several times as described in the hydrogen peroxide/osmotic stress assays section above. However, cultures were placed in a 30°C shaker instead of a 30°C rolling incubator. Exponential phase cultures were harvested when the last starter culture (i.e., the OD_660_=0.05 culture at the 16 hour time point) reached an OD_660_ of 0.2-0.25. Saturated growth cultures were harvested when the exponential phase culture reached an OD660 of 0.85-0.90. Reporter assays in which the effect of stress treatment was quantified were conducted on exponential phase cultures that were split immediately before treatment.

β-galactosidase activity from chloroform-permeabilized cells was measured using the colorimetric substrate o-nitrophenyl-β-D-galactopyranoside (ONPG). 1 mL enzymatic reactions contained 200-250 μL of chloroform-permeabilized cells, 550-600 μL of Z-buffer (60 mM Na_2_HPO_4_, 40 mM NaH_2_PO_4_, 10 mM KCl, 1 mM MgSO_4_), and 200 μL of 4 mg/mL ONPG in 0.1 M KPO4, pH 7.0. Chloroform-permeabilized cell samples were prepared from 100-150 μL of culture, 100 μL of PYE, and 50 μL of chloroform (chloroform volume is not included in the final calculation of the 1 mL reaction). Chloroform-treated cells were vortexed for 5-10 seconds to facilitate permeabilization. Z buffer and ONPG were added directly to chloroform-permeabilized cells. Reactions were incubated in the dark at room temperature and quenched with 1 mL of 1 M Na_2_CO_3_.

Each reporter construct was optimized with different reaction times and different volumes of cells. Reaction time and volume for each reporter was empirically determined by the development of the yellow pigment from chloroform-permeabilized *C. crescentus* CB15 cultures. Strains harboring the pRKlac290 transcriptional reporter plasmid containing the established GSR promoter reporter P*_sigU_* or P*_gsrN_* used 100 μL of cells and were quenched after 10 minutes and 18 minutes, respectively. Strains containing pRKlac290 with the *katG* promoter (P*_katG_*) used 150 μL of cells and were quenched after 12 minutes. Strains with the translational reporter plasmid pPR9TT containing the 5’UTR of *katG* (wildtype and RS constructs) used 150 μL of cells and were quenched after 4 minutes.

Miller units were calculated as:

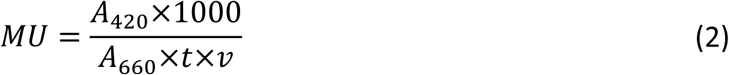

Where A_420_ is the absorbance of the quenched reaction measured at 420 nm on a Spectronic Genesys 20 spectrophotometer (ThermoFisher Scientific, Waltham, MA). A_660_ is the optical density of the culture of cells used for the assay. *t* is time in minutes between the addition of ONPG to the time of quenching with Na_2_CO_3_. *𝑣* is the volume in milliliters of the culture added to the reaction.

#### TRIzol RNA extractions

Cultures used for the extraction of RNA were passaged in the same manner outlined in the hydrogen peroxide/osmotic stress assays section above. Exponential phase cultures were harvested from the last starter (i.e., the OD_660_=0.05 culture at the 16 hour time point) when it reached an OD_660_ of 0.20-0.25. Saturated cultures were harvested when the final culture diluted to OD_660_=0.025 reached an OD_660_ of 0.85-0.90.

Exponential phase cultures (OD_660_ of 0.20-0.25) harvested for extraction of RNA were pelleted at 15000xg for 3 minutes at ≈23°C (i.e. room temperature). Early saturated cultures (OD_660_ of 0.85-0.90) were also pelleted at 15000xg for 30 seconds at ≈23°C. All media were aspirated using a vacuum flask. Cell pellets were resuspended in 1 mL of TRIzol^™^. The TRIzol resuspension was heated for 10 minutes at 65°C, treated with 200 μL of chloroform and hand shaken. The chloroform mixture was allowed to stand for 5 minutes and then spun down at 15000xg for 15 minutes at 4°C. Approximately 500 μL of clear aqueous phase was extracted and mixed with 500 μL of 100% isopropanol. Samples were then incubated at −20°C overnight. Overnight isopropanol precipitation was then spun down at 15000xg for 15 minutes at 4°C. Isopropanol was aspirated, the pellet was washed in 1mL of 75% ethanol, and sample was spun down at 15000xg for 15 minutes at 4°C. Ethanol was removed from pellet, and the pellet was left to dry for 15 minutes. The RNA pellet was resuspended in 25 μL of nuclease-free H_2_O.

#### Radiolabeled Oligonucleotides

Oligonucleotides were radiolabeled with T4 Polynucleotide Kinase (PNK). 10μL labeling reactions were composed of 1μL of PNK, 1μL PNK 10× Buffer, 2μL of 5 μM oligonucleotides (1 μM final concentration), 4μL H_2_O, and 2 μL ATP, [γ-32P]. Reactions were incubated for a minimum of 37°C for 30 minutes. Total reactions were loaded onto a BioRad P-6 column to clean the reaction. Radiolabeled samples were stored at 4°C.

#### Northern Blots

RNA samples were resolved on a 10% acrylamide:bisacrylamide (29:1), 7 M urea, 89 mM Tris Borate pH 8.3, 2 mM Na_2_EDTA (TBE) 17 by 15 cm gel, run for 1 hour and 50 minutes at 12 Watts constant power in TBE running buffer. The amount of sample loaded was between 1-5 μg of RNA, mixed in a 1:1 ratio with 2x RNA loading dye (9 M urea, 100 mM EDTA, 0.02% w/v xylene cyanol, 0.02% w/v bromophenol blue). Samples were heated for 8 minutes at 75°C and then subjected to an ice bath for 1 minute before loading. Acrylamide gels with immobilized samples were then soaked in TBE buffer with ethidium bromide and imaged. Samples immobilized on the gel were transferred onto ZetaProbe Blotting Membrane with a Trans-Blot^®^ SD Semi-Dry Transfer Cell. Transfer was done at 400 mA constant current with voltage not exceeding 25V for 2 hours. Membrane was then subjected to two doses of 120 mJ/cm^2^ UV radiation, using a Stratalinker UV cross-linker. Membranes were subsequently prehybridized 2 times for 30 minutes in hybridization buffer at 65°C in a rotating hybridization oven. Hybridization buffer is a variation of the Church and Gilbert hybridization buffer (20 mM sodium phosphate, pH 7, 300 mM NaCl, 1% SDS). Blots were hybridized with hybridization buffer containing the radiolabeled oligonucleotide probes described above. Hybridization buffer was always prepared so that GsrN probe concentration was approximately 1 nM, 5S rRNA probe concentration was approximately 2 pM, and tRNA-Tyr probe was 500 pM. Hybridization took place over 16 hours at 65°C in a rotating hybridization oven. Membranes were then incubated with wash buffer three times for 20 minutes at 65°C in a rotating hybridization oven. Wash buffer contained 20 mM sodium phosphate (pH 7.2), 300 mM NaCl, 2 mM EDTA, and 0.1% SDS. Membranes were then wrapped in plastic wrap and placed directly against a Molecular Dynamics Phosphor Screen. Screens were imaged with Personal Molecular Imager™ (PMI™) System. Membrane exposure time was determined using a Geiger counter: 100× 2 minutes, 10× 30-60 minutes, 1.0× 8-16 hours, 0.1× 48-72 hours.

Intensity of GsrN bands or *katG* mRNA dots was calculated by dividing the probe signal specific to GsrN or *katG* mRNA over the probe signal specific to the 5S rRNA multiplied by 100. Normalization of *katG* mRNA specific probes in the dot blot was carried out in a manner similar to that described for Northern blot, in which the 5S rRNA probe signal was used for normalization.

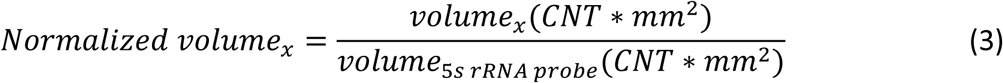

#### Rifampicin transcription inhibition assays

Liquid *C. crescentus* CB15 cultures were passaged in the same manner outlined in the hydrogen peroxide/osmotic stress assays section. However, cells for transcription inhibition assays were grown to an OD_660_ of 0.2-0.25 from the last starter culture (i.e., inoculated from the OD_660_=0.05 culture from 16 hour growth) and split across 6 tubes and labeled: untreated, 30 second treatment, 2 minute treatment, 4 minute treatment, 8 minute treatment, and 16 minute treatment. Untreated cultures were the 0 time point where no rifampicin was added. Rifampicin treated cultures were subjected to a final concentration of 10 μg/mL (from a 10 mg/mL stock in methanol) and were grown in a rolling incubator at 30°C. The 30 second rifampicin treatment refers to the centrifugation time (15000xg for 30 seconds at room temperature) to pellet the cells. Thus, the 30 second sample was immediately pelleted after exposure to rifampicin. 2 minute, 4 minute, 8 minute, and 16 minute samples were placed into a rolling incubator after exposure and were removed 30 seconds prior to their indicated time point, (i.e. 2 minute culture was removed from the incubator at 1 minute and 30 seconds). Pellets were then subjected to TRIzol extraction as described earlier. RNA extracts were subjected to Northern Blot analysis as described earlier.

Intensity of full-length and 5’isoform of GsrN bands were first adjusted to the intensity of the 5S rRNA control, as described in Equation 3. To plot the GsrN decay curve, all adjusted bands were then divided by the intensity of the 0 time point (untreated culture) and plotted in Prism v6.04.

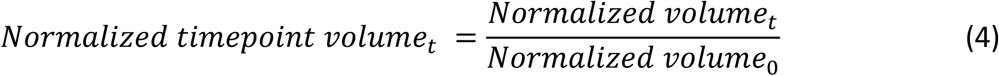

#### Primer extension

Primer extension was carried out using the SuperScript™ IV Reverse Transcriptase standalone enzyme. Total RNA from *gsrN^++^* and Δ*gsrN* strains was extracted from saturated cultures (OD_660_=0.95-1.0) as described in the TRIzol extraction section. Primers for extension were first HPLC purified (Integrated DNA technologies) and radiolabeled as described in the Radiolabeled Oligonucleotides section.

Briefly, 14 μL annealing reactions comprised of the following final concentrations/amounts: 0.1 μM of gene specific radiolabeled primer, 0.3-0.5 mM of dNTPs, 2 μg of total RNA, and when necessary 0.5 mM ddNTPs. ddNTP reactions had a 3 dNTP:5 ddNTP ratio and were conducted using total RNA from *gsrN^++^*. Annealing reactions were incubated at 65°C for 5 minutes and subsequently incubated on ice for at least 1 minute.

Extension reactions contained 14 μL annealing reactions with 6 μL of SuperScript™ IV Reverse Transcriptase master mix (final concentrations/amount 5 mM DTT, 2.0 U/μL, 1x SSIV buffer). Reactions were incubated at 50–55°C for 10 minutes and then incubated at 80°C for 10 minutes to inactivate the reaction.

After the extension reaction, 1 μL of RNase H was added to the mixture. This was incubated at 37°C for 20 minutes and mixed with 20 μL of 2x RNA loading dye. Reactions were subsequently heated for 8 minutes at 80°C, subjected to an ice bath for 1 minute, and loaded onto a 33.8 by 19.7 cm 20% acrylamide:bisacrylamide gel (as outlined in the Northern Blot section). Reactions were loaded on the gel along with a labeled Low Molecular Weight Marker (10-100 nt; Affymetrix/USB). Final amounts loaded were estimated using a Geiger counter, such that 10 mR/hr was loaded for each sample. Primer extension samples were resolved on the gel at 10 Watts constant power until unextended primer reached the bottom of the gel. The acrylamide gel was wrapped in plastic, exposed, and imaged as outlined in the Northern Blot section.

#### Affinity purification of GsrN using a PP7hp-PP7cp system

GsrN constructs containing a Pseudomonas phage 7 RNA hairpin (PP7hp) sequence were affinity purified using a hairpin-binding phage coat protein (PP7cp) immobilized on agarose beads. To prepare the coat protein, a 50 mL culture of *E. coli* Rosetta carrying an expression plasmid for PP7cp fused to maltose binding protein (MBP) at its N-terminus and a His-tag at its C-terminus (pMal-PP7cp-HIS) was grown at 37°C in a shaking incubator overnight in LB-ampicillin broth. Overnight cultures were rediluted and grown to OD_600_=0.6. Cells were then induced with 1mM IPTG for 5 hours and spun down at 8000g at 4°C for 10 minutes. The cell pellet was resuspended in 6 mL of ice-cold lysis buffer (125 mM NaCl, 25 mM Tris-HCl pH 7.5, 10 mM Imidazole) and mechanically lysed in a LV1 Microfluidizer. Lysate was immediately added to 500 μL of amylose resin slurry that was prewashed with ice-cold lysis buffer. After the sample was loaded, beads were washed in 50x bead volume (^~^10mL) of ice-cold lysis buffer.

A 50 mL culture of *C. crescentus* Δ*gsrN* carrying plasmid pMT552 expressing PP7hp-tagged alleles of *gsrN* was grown at 30°C in a shaking incubator overnight in M2X medium. The culture was prepared from a starter and passaged as outlined in the hydrogen peroxide/osmotic stress assays section. Cells were grown to an OD_660_=0.85-0.90. Cells were spun down at 8000g at 4°C for 15 minutes, resuspended in 6 mL of ice-cold lysis buffer, and mechanically lysed in a LV1 Microfluidizer. Lysate was immediately loaded onto a column of amylose resin on which pMal-PP7cp-HIS had been immobilized. After the sample was loaded, beads were washed in 50x bead volume (^~^10mL) of ice-cold lysis buffer. Elution of MBP-PP7cp-HIS bound to GsrN-PP7hp and associated biomolecules was completed over three 0.5 mL elution steps using 500 mM maltose. Each 0.5 mL elution was then mixed with equal volumes of acid-phenol for RNA extraction for RNA analysis, or equal volumes of SDS-Loading Buffer (200 mM Tris-HCl pH 6.8, 400 mM DTT, 8% SDS, 0.4% bromophenol blue, 40% glycerol) for protein analysis. For the RNA analysis, the three elution fractions were combined in an isopropanol precipitation step. RNA samples were subjected to DNase treatment as outlined in the RNA-seq sequencing section.

#### Acid-Phenol RNA extraction

Samples for acid-phenol extractions were mixed with equal volumes of acid-phenol and vortexed intermittently at room temperature for 10 minutes. Phenol mixture was spun down for 15 minutes at maximum speed at 4°C. The aqueous phase was extracted, cleaned with an equal volume of chloroform, and spun down for 15 minutes at maximum speed at 4°C. The aqueous phase was extracted from the organic and equal volumes of 100% isopropanol were added. Linear acrylamide was added to the isopropanol precipitation to improve pelleting (1 μL per 100 μL of isopropanol sample). Samples were then incubated at −20°C overnight and spun down at 15000xg for 15 minutes at 4°C. The isopropanol was aspirated, the pellet washed in 1 mL of 75% ethanol, and sample spun again at 15000xg for 15 minutes at 4°C. Ethanol was removed from the RNA pellet, and pellet was left to dry for 15 minutes. Pellet was resuspended in 25 μL of nuclease-free H_2_O.

#### RNA dot blot analysis

Samples (≈3 μg) for dot blot analysis were mixed with equal volumes of 2x RNA loading dye as in a Northern Blot, and heated for 8 minutes at 75°C. Samples were then spotted on a Zeta-Probe Blotting Membrane and left to dry for 30 minutes. Spotted membrane was then subjected to two doses of 120 mJ/cm^2^ UV radiation (Stratalinker UV crosslinker). The membrane was then prehybridized 2 times for 30 minutes in hybridization buffer at 65°C in a rotating hybridization oven. After prehybridization, we added radiolabeled oligonucleotide probes. Hybridization buffer with probes was always prepared so that each probe’s concentration was approximately 1 nM. *katG* mRNA was first hybridized for 16 hours at 65°C in a rotating hybridization oven. Membrane was then washed with wash buffer three times, 20 minutes each at 65°C in a rotating hybridization oven. The blot was exposed for 48 hours to a Molecular Dynamics Phosphor screen and imaged on a Personal Molecular Imager as described above. Membrane was subsequently stripped with two rounds of boiling in 0.1% SDS solution and incubated for 30 minutes at 65°C in a rotating hybridization oven. Following stripping, the membrane was subjected to two rounds of prehybridization and then hybridized for 16 hours at 65°C in a rotating hybridization oven with the probe specific to the 5’ end of GsrN. Membrane was then washed again with wash buffer three times for 20 minutes each at 65°C in a rotating hybridization oven. This GsrN blot was exposed for 36 hours to the phosphor screen and imaged. The membrane was stripped four times after GsrN probe exposure. Following stripping, membrane was again subjected to two rounds of prehybridization and then hybridized for 16 hours at 65°C in a rotating hybridization oven with the probe specific to 5S rRNA. Membrane washed with Wash Buffer three times, 20 minutes each at 65°C in a rotating hybridization oven. This 5S RNA blot was exposed to the phosphor screen for 1 hour and imaged.

#### Western Blot analysis

Strains from which protein samples were prepared for Western blot analysis were grown and passaged as outlined in the hydrogen peroxide/osmotic stress assays section. However, cultures were taken from the overnight 16-hour growth when OD_660_ reached 0.85-0.90. 1 mL of these cultures was then pelleted, resuspended in 125 μL of Western blot buffer (10 mM Tris pH 7.4, 1 mM CaCl_2_, and 5 μg/mL of DNase), and mixed with 125 μL SDS-Loading buffer. Samples were boiled at 85°C for 10 minutes, and 10-20 μL of each sample was loaded onto a Mini-PROTEAN TGX Precast Gradient Gel (420%) with Precision Plus Protein™ Kaleidoscope™ Prestained Protein Standards. Samples were resolved at 35 mA constant current in SDS running buffer (0.3% Tris, 18.8% Glycine, 0.1% SDS). Gels were run until the 25 kDa marker reached the bottom of the gel. Gel was transferred to an Immobilon^®^-P PVDF Membrane using a Mini Trans-Blot^®^ Cell after preincubation in Western transfer buffer (0.3% Tris, 18.8% Glycine, 20% methanol). Transfer was carried out at 4°C, 100 V for 1 hour and 20 minutes in Western transfer buffer. The membrane was then blocked in 5% (w/v) powdered milk in Tris-buffered Saline Tween (TBST: 137 mM NaCl, 2.3 mM KCl, 20 mM Tris pH 7.4, 0.1% (v/v) of Tween 20) overnight at room temperature on a rotating platform. Primary incubation with a DYKDDDDK(i.e. M2)-Tag Monoclonal Antibody (clone FG4R) was carried out for 3 hours in 5% powdered milk TBST at room temperature on a rotating platform (4 μL antibody in 12 mL). Membrane was then washed 3 times in TBST for 15 minutes each at room temperature on a rotating platform. Secondary incubation with Goat anti-Mouse IgG (H+L) Secondary Antibody, HRP was for 1 hour at room temperature on a rotating platform (3 μL antibody in 15 mL). Finally, membrane was washed 3 times in TBST for 15 minutes each at room temperature on a rotating platform. Chemiluminescence was performed using the SuperSignal™ West Femto Maximum Sensitivity Substrate and was imaged using a ChemiDoc MP Imaging System version 6.0. Chemiluminescence was measured using the ChemSens program with an exposure time of ^~^2 minutes.

Western blot lane normalization of KatG-M2 specific bands was conducted by normalizing total signal from the doublet signal in the M2 specific background to that of the non-specific band (found in strains were there was no M2 tagged KatG). Samples extracted on the same day were run on the same gel. Lane normalized samples were then normalized to the levels of KatG-M2 signal in the wildtype untreated samples for that specific gel.

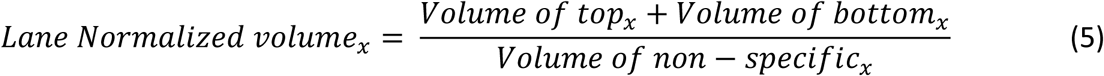

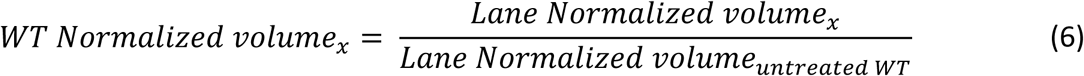

#### RNA-seq preparation

Total RNA was extracted from cultures passaged similarly to the hydrogen peroxide/osmotic stress assays section. However, cultures were harvested at OD_660_=0.85-0.90 from the 16-hour overnight growth. Total RNA extraction followed the procedure outlined in the TRIzol extraction section. Resuspended RNA pellets after the 75% ethanol wash were loaded onto an RNeasy Mini Kit column (100 μL sample, 350 μL RLT, 250 μL 100% ethanol). Immobilized RNA was then subjected to an on-column DNase digestion with TURBO™ DNase. DNase treatment was repeated twice on the same column; each incubation was 30 minutes at 30°C with 70 μL solutions of DNase Turbo (7 μL DNase, 7 μL 10x Buffer, 56 μL diH_2_O). RNA was eluted from column, rRNA was depleted using Ribo-Zero rRNA Removal (Gram-negative bacteria) Kit (Epicentre). RNA-seq libraries were prepared with an Illumina TruSeq stranded RNA kit according to manufacturer’s instructions. The libraries were sequenced on an Illumina HiSeq 4000 at the University of Chicago Functional Genomics Facility.

#### Soluble protein extraction for LC-MS/MS proteomics

Total soluble protein for proteomic measurements was extracted from cultures passaged similarly to the hydrogen peroxide/osmotic stress assays section. However, harvested cultures were grown to an OD_660_=0.85-0.90 in 50 mL of M2X during the 16-hour overnight growth in a 30°C shaking incubator. Cells were spun down at 8000g at 4°C for 15 minutes. Cells were resuspended in 6 mL of ice-cold lysis buffer. Cells were mechanically lysed in LV1 Microfluidizer. Lysate was then spun down at 8000g at 4°C for 15 minutes. Protein samples were resolved on a 12% MOPS buffered 1D Gel (Thermo Scientific) for 10 minutes at 200V constant. Gel was stained with Imperial Protein stain (Thermo Scientific), and a ^~^2 cm plug was digested with trypsin. Detailed trypsin digestion and peptide extraction by the facility is published in (Truman et al., 2012).

#### LC-MS/MS data collection and analysis

Samples for analysis were run on an electrospray tandem mass spectrometer (Thermo Q-Exactive Orbitrap), using a 70,000 RP survey scan in profile mode, m/z 360-2000 Fa, with lockmasses, followed by 20 MSMS HCD fragmentation scans at 17,500 resolution on doubly and triply charged precursors. Single charged ions were excluded, and ions selected for MS/MS were placed on an exclusion list for 60s (Truman et al., 2012).

### Computational Methods

#### Network construction

RNAseq data (15 read files) was obtained from the NCBI GEO database from (Fang et al., 2013). Read files are comprised of 3 biological replicates of total RNA extracted from *C. crescentus* cultures at 5 time points across the cell cycle (0, 30, 60, 90, and 120 minutes post synchrony). Reads were mapped and quantified with Rockhopper 2.0 (Tjaden, 2015). The estimated expression levels of each gene across the 5 time points were extracted from the “Expression” column in the “_transcripts.txt” file, using the “verbose” output. Expression of each gene across the 5 time points was normalized using python scripts as follows: for a given gene, the normalized expression of the gene at a time point, t, is divided by the sum of the gene’s expression across all the time points, Equation 7. Thus the sum of a gene’s normalized expression across the 5 time points would equal 1.

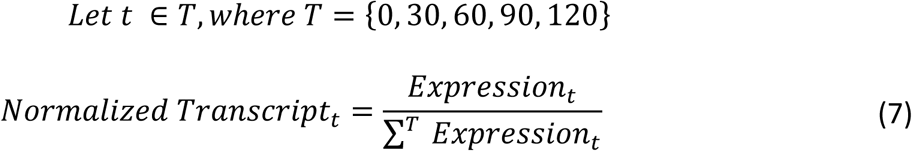

We computed Pearson’s correlation coefficient based on normalized expression between all pairwise combinations of genes. Correlation coefficients were organized into a numpy.matrix data structure where each row and column corresponds to the same gene order. Correlation coefficients less than 0 were not considered for this analysis and were assigned the value 0. We refer to this matrix as the Rho-matrix. The Rho-matrix is symmetric and the product of its diagonal is 1. The Rho-matrix represents the weighted edges of the network, where the value of 0 demonstrates no edge is drawn between nodes.

A one-dimensional weight matrix that corresponds to the rows and columns of the Rho-matrix was constructed as a numpy.matrix data structure with all values initialized at 0. Lastly, a key array was constructed in conjunction with the Rho-matrix and weight-matrix for initializing the assignment of weight and obtaining the final weights of the algorithm. The weight-matrix represents the weight of the nodes of the network and the key matrix represents the gene name of the node.

#### Iterative Ranking: Matrices and Algorithms

Iterative ranking algorithms are a class of analytical tools used to understand relationships between nodes of a given network. The iterative ranking algorithm used to dissect the general stress response in the transcription-based network follows:

Given the Rho-matrix (P) and weight-matrix (*f*), the weight-matrix after *t*-iterations is Equation 8.

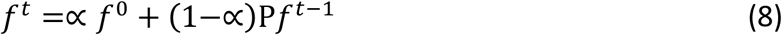

For Equation 8, let *∝* represent a dampening factor applied to the initialize (*t*) = 0 weight of the nodes, *f*^0^. The final weights of the weight-matrix as *t* → ∞ converge to a stable solution, Equation 9.

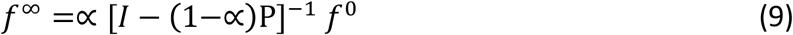

Algorithm and solution information was adapted from (Wang and Marcotte, 2010).

Initial weight-matrix, (*f*^0^), was created by assigning the weight 1.0 to the corresponding positions of the seven genes known to regulate the General Stress Response (GSR) of *C. crescentus*: *sigT*, *phyR*, *phyK*, *sigU*, *nepR*, *lovR*, and *lovK*. Normalization of the values of the Rho-matrix, P, was performed by normalizing each column such that each column has a sum equal to 1 and then repeating the same normalization process by rows.

#### Iterative rank parameter tuning

Iterative rank parameters were optimized through the self-prediction of known associated components of the General Stress Response (GSR). Variables tuned for exploration were the *∝* parameter and the reduction of the number of edges based on correlation cut-offs. We chose to base our parameters on which condition best predicted the gene *phyR*, when initializing the weight-matrix with *sigT*,*sigU*, *nepR*, *phyK*, *lovR*, and *lovK* values of 1. Varying these two parameters showed that an edge reduction of ρ > 0.9 and an alpha factor greater than 0.5 yielded the highest rank for *phyR* (Figure 1- figure supplement 1A).

A ρ > 0.9 edge reduction reduces the number of edges each node has (Figure 1- figure supplement 1B). The total number of edges was reduced from 10225998 edges to 946558 (Figure 1- figure supplement 1C). Only 19 nodes (.46%) were completely disconnected from the network (zero number of edges). Tuning script is available at https://github.com/mtien/IterativeRank.

#### Identification of σ^T^-promoter motifs

Motif finder utilized a python script that scans 200 nucleotides upstream of annotated transcriptional start sites (Zhou et al., 2015) or predicted translational start sites (TSS) (Marks et al., 2010).

We built a simple python library to take in genomic FASTA files, find specified regions of interest, and extract 200 nucleotides from a given strand. We used the *Caulobacter crescentus* NA1000 annotation (CP001340) from NCBI as the input genomic file and used the predicted TSS (when available) or annotated gene start sites as the region and strand specifier. After locating the position and strand within the file, we extracted the 200 nucleotides directly upstream of the site of interest and put the regions into a character-match calculator. Our simple calculator reported a list of positions for −35 (GGAAC) and for −10 elements (CGTT) of σ^T^-dependent promoters within the 200-nucleotide input string. Only strict matches to these elements were reported. Spacers were calculated between all pairwise −35 and −10 matches. We identified potential σ^T^-dependent promoters by identifying consensus −35 to −10 sequences with 15-17 base spacing. Sequence logos were generated from (Crooks et al., 2004)

#### IntaRNA analysis

IntaRNA version 2.0.2 is a program within the Freiberg RNA Tools collection (Mann et al., 2017). To predict likely RNA-RNA associations between predicted unstructured regions within GsrN and its RNA targets, we input the sequence of GsrN as the query ncRNA sequence and a FASTA file of either: 1) windows significantly enriched in the GsrN(37)-PP7hp purification from our sliding window analysis with an additional 100 base pairs (50 bp on each side of the window) or 2) entire gene windows that showed significant enrichment from our Rockhopper analysis (Figure 5 – source table 3).

Output from IntaRNA comprised a csv file of target binding sites and the corresponding GsrN binding sites. We extract the predicted binding sites of the targets with a python script and parsed the targets into those predicted to bind the first exposed loop and the second exposed loop. Sequence logos were generated from (Crooks et al., 2004)

#### Phylogenetic tree construction

A 16S rRNA phylogenetic tree of Alphaproteobacteria was constructed by extracting 16S rRNA sequences for all species listed in Figure S7A and using the tree building package in Geneious 11.0.2 (Kearse et al., 2012). The tree was constructed using a global alignment with free end gaps and a cost matrix of 65% similarity (5.0/^~^4.0). The genetic distance model was the Tamura-Nei and the tree building method employed was neighbor-joining. *E. litoralis* was the out-group for tree construction.

#### Prediction of *gsrN* homologs

A homology search based on the sequence of GsrN was conducted using BLASTn (Altschul et al., 1990). This simple search provided a list of clear GsrN homologs in the Caulobacteraceae family (*Caulobacter*, *Brevundimonas*, and *Phenylobacterium*).

Identification of homologs in other genera relied on analysis of published transcriptomic data, searching specifically for gene expression from intergenic regions. Analyzed data included *Rhizobium etli* (Jans et al., 2013), *Sinorhizobium meliloti* (Valverde et al., 2008) and *Brucella abortus* (Kim et al. 2014). The prediction of GsrN homologs in *Rhodopseudomonas palustris* and *Bradyrhizobium diazoefficiens* is completely based on the proximity of a GsrN-like sequence to the GSR locus and the presence of a σ^ecfG^ site in the predicted promoters of these predicted genes.

#### Mapping reads from RNA-seq data

RNA-seq read files (fastQ) were aligned with sequence files (fastA) using bowtie 2.0 (Langmead and Salzberg, 2012). SAMTools was then used to calculate the depth and coverage of each nucleotide in the hit output file from bowtie 2.0 (Li et al., 2009). Normalization of reads per nucleotide was computed by normalizing each count to the per million total number of reads mapped to all of the CP001340.1 genome. Normalized reads per nucleotide was then plotted in Prism v6.04 where standard error and mean were calculated.

#### RNA-seq analysis of mRNAs that co-elute with GsrN

RNA-seq read files (fastQ) from the three replicate GsrN(37)⸬PP7hp purifications and duplicate PP7hp-GsrN-3’ purifications were quantified and analyzed with Rockhopper 2.0 (Tjaden, 2015). Reads were mapped to modified *C. crescentus* genome files (fastA, PTT, RNT) where the wild-type *gsrN* locus was replaced with the sequence of *gsrN(37)-PP7hp.* Using the “verbose output” option and the resulting “transcripts.txt” file, we pruned the dataset to find genes that had low FDR values (“qValue” <.05), were significantly enriched in GsrN(37)⸬PP7 (“ Expression GsrN(37)-PP7hp” > “Expression PP7hp-GsrN-3’”), and had a high total number of reads that mapped to GsrN(37)⸬PP7 (“Expression GsrN(37)-PP7hp” >1000). This analysis provided a list of 35 candidate genes (Figure 5-source data 1).

The Rockhopper analysis package organizes reads into IGV (integrative Genomic Viewer) files. Upon visual inspection and spot validation of the 35 candidates in IGV, we found 26 genes with consistently higher signal across the three GsrN(37)⸬PP7hp purifications relative to PP7hp-GsrN-3’ control fractions. In some cases, reads mapped outside coding sequences. Such reads mapped proximal to the 5’ end of annotated genes and to intergenic regions. We observed uneven read distribution across some annotated genes. Cases in which reads were not evenly distributed across a gene were typically not classified as significantly different from the control samples in “Expression” or “qValue” by Rockhopper even when a clear bias in read density was visually evident (most often at the 5’ end of the gene).

As a second approach, we performed a systematic window annotation analysis to capture the unaccounted read density differences between the two purified fractions (GsrN(37)⸬PP7hp and the PP7hp-GsrN-3’ negative control). Windows were generated by *in silico* fragmentation of the *C. crescentus* NA1000 genome sequence, designating 25 base pair windows across the genome. We prepared new annotated window files (FASTA, PTT, RNT) for wild-type, *gsrN(37)*-*PP7hp*, and *PP7hp*-*gsrN*-3*’*. The window identification number corresponds to the same sequence across the three different FASTA sequences.

Mapping and quantification of reads to these windows was conducted using the EDGE-pro analysis pipeline (Magoc et al., 2013). A caveat of EDGE-pro quantification is the potential misattribution of reads to input windows. EDGE-pro quantification does not take strand information into account when mapping reads to input windows.

Read quantification of the *gsrN(37)⸬PP7hp* purifications showed consistent differences in one of the three samples. *gsrN(37)⸬PP7hp* sample 1 contained 2.69% reads mapped to *gsrN(37)*-*PP7hp* while sample 2 and 3 had 15.78% and 14.04% mapped to *gsrN(37)*-*PP7hp* respectively. Additionally, we observed that sample 1 had several genes that were strongly enriched in sample 1 and not in sample 2 and 3. Thus we employed a metric to balance the discrepancies between the three separate purifications. To minimize potential false positives, we calculated the average of all three samples and the average of samples 2 and 3. If the total average was 1.5 times greater than the sample 2 and 3 average, we assumed that the sample 1 artificially raised the average RPKM value and did not consider any data from any of the purifications in that specific window. The total window population decreased from 161713 windows to 109648 windows after this correction. This process is reflected in the https://github.com/mtien/Slidingwindowanalysis script “remove_high_variant_windows.py”.

From the RPKM values calculated with EDGE-pro, we used the R-package, DESeq (Anders and Huber, 2010), to assess statistically significant differences between windows of expression. Candidate windows enriched in the GsrN(37)⸬PP7 fractions were identified using metrics similar to what is applied to traditional RNA-seq data. Briefly, we identified windows that had a low p-values ( pvalue< .10), were enriched in the GsrN(37)⸬PP7 (“baseMean GsrN(37)-PP7hp” > “baseMean PP7hp”), and had a high level of reads mapped to the gene in the GsrN(37)⸬PP7 (“baseMean GsrN(37)-PP7hp” >1000) (Figure 5-source data 2). Since the read density of windows from the total RNA extracted from the PP7-purification did not converge when estimating dispersion with a general linear model, we added total RNA seq read density from wild-type strains grown in stationary phase to help model the dispersion for the negative binomial analysis by DESeq, GSE106168.

Adjacent significant windows were then combined and mapped onto the annotated genome of *C. crescentus.* In order to correct for strand information lost in EDGE-pro quantitation, bowtie file information was used to define the strand of reads mapped to combined significant windows (Table 1).

#### RNA-seq processing of total RNA

Analysis of whole genome RNA-seq data was conducted using the CLC Genomics Workbench (Qiagen). Reads were mapped to the *C. crescentus* NA1000 genome (accession CP001340.1) (Marks et al., 2010). Differential expression was determined using Wald test in the CLC Workbench suite (Figure 8- source table 2).

#### LC-MS/MS processing of total soluble protein

Raw files of LC-MS/MS data collected on wild-type, Δ*gsrN*, and *gsrN^++^* were processed using the MaxQuant software suitev1.5.1.2 (Cox et al., 2014). Samples were run against a FASTA file of proteins from the UniProt database (UP000001364) and standard contaminants. The label free quantitation (LFQ) option was turned on. Fixed modification included carbamidomethyl (C) and variable modifications were acetyl or formyl (N-term) and oxidation (M). Protein group files were created for three comparisons: wild-type versus Δ*gsrN*, Δ*gsrN* versus *gsrN^++^*, and wild-type versus *gsrN^++^* samples.

LFQ values for each protein group were compiled across all three runs and used as estimated protein quantities in our analyses (Figure 8A). Each strain had a total of 6 LFQ values for every protein group, 2 from each of the comparisons. Average LFQ values were only calculated if 3 or more LFQ values were found for a given protein group. This allowed for protein groups that had a sufficient amount of signal across all the samples and analyses to be considered for comparison. Once averages for each protein group were calculated, we calculated the fold change between samples from different backgrounds by dividing the averages and taking the log-2 transformation (log2Fold).

Multiple t-tests were conducted using all 6 LFQ value obtained across the three MaxQuant runs. We used the multiple t-test analysis from GraphPad Prism version 6.04 for MacOS, GraphPad Software, La Jolla California USA, www.graphpad.com. The false discovery rate (Q) value was set to 5.000% and each row was analyzed individually, without assuming a consistent SD.

#### Data And Software Availability

IterativeRank and RhoNetwork python libraries are available on https://github.com/mtien/IterativeRank.

Scripts used to analyze the total RNA reads from the PP7-affinity purification are available on https://github.com/mtien/Sliding_window_analysis.

RNA-seq data of wild-type, Δ*gsrN*, and *gsrN^++^* in early stationary cultures are deposited in the NCBI GEO database under the accession number GSE106168.

RNA-seq affinity purification data have been deposited in the NCBI GEO database under accession number GSE106171.

LC-MS/MS data is available on the PRIDE Archive EMBL-EBI under the accession number PXD008128.

Key Reagent And Resource Table

**Key Reagent And Resource Table**
| REAGENT or RESOURCE | SOURCE | IDENTIFIER |
| --- | --- | --- |
| **Antibodies** |  |  |
| Goat anti-Mouse IgG (H+L) Secondary Antibody, HRP | ThermoFisher | 32430 |
| DYKDDDDK Tag Monoclonal Antibody (FG4R) | ThermoFisher | MA1-91878-1MG |
| **Bacterial and Virus Strains** |  |  |
| See Supplementary File 1 |  |  |
| **Chemicals, Peptides, and Recombinant Proteins** |  |  |
| Agar | Lab Scientific | A466 |
| 30% Hydrogen Peroxide | ThermoFisher | H325-100 |
| substrate o-nitrophenyl-β-D-galactopyranoside (ONPG) | GoldBio | N-275-100 |
| acrylamide:bisacrylamide (29:1) | BioRad | 1610156 |
| Acid-Phenol | Ambion | Am9722 |
| TRIzol | ThermoFisher | 15596026 |
| T4 Polynucleotide Kinase | New England Biolabs | M0201L |
| ATP, [γ-32P]- 3000Ci/mmol 10mCi/ml EasyTide | PerkinElmer | BLU502A500UC |
| SuperScript™ IV Reverse Transcriptase | ThermoFisher | 18090010 |
| RNase H | New England Biolabs | M0297S |
| TURBO™ DNase | ThermoFisher | AM2238 |
| **Critical Commercial Assays** |  |  |
| Micro Bio-Spin Columns With Bio-Gel P-6 in Tris Buffer | BioRad | 7326221 |
| Amylose Resin | New England Biolabs | E8021L |
| RNeasy Mini Kit | Qiagen | 74106 |
| SuperSignal™ West Femto Maximum Sensitivity Substrate | ThermoFisher | 34095 |
| **Deposited Data** |  |  |
| Raw and analyzed RNA-seq data | This paper | GEO: GSE106168<br>https://www.ncbi.nlm.nih.gov/geo/query/acc.cgi?acc=GSE106168 |
| Raw and analyzed LC-MS/MS data | This paper | PRIDE: PXD008128 |
| Raw and analyzed RNA-seq data for GsrN-PP7hp purification | This paper | GEO: GSE106171<br>https://www.ncbi.nlm.nih.gov/geo/query/acc.cgi?acc=GSE106171 |
| Raw and analyzed RNA-seq data for Network construction | (Fang et al., 2013) | GEO: GSE46915 |
| **Gel and Blotting equipment** |  |  |
| Zeta-Probe Blotting Membranes | BioRad | 162-0165 |
| Low Molecular Weight Marker, 10-100 nt | Alfa Aesar | J76410 |
| Mini-PROTEAN TGX Precast Gel, 4-20% | BioRad | 456-1094 |
| Precision Plus Protein™ Kaleidoscope™<br>Prestained Protein Standards | BioRad | 1610375 |
| Oligonucleotides |  |  |
| See Supplementary File 1 |  |  |
| Plasmids |  |  |
| See Supplementary File 1 |  |  |
| Software and Algorithms |  |  |
| Bowtie2 | (Langmead and Salzberg, 2012) | http://bowtie-bio.sourceforge.net/bowtie2/index.shtml |
| SAMTools | (Li et al., 2009) | http://samtools.sourceforge.net/ |
| IntaRNA | (Mann et al., 2017) | http://rna.informatik.uni-freiburg.de/IntaRNA/Input.jsp |
| Prism v6.04 | GraphPad Software, Inc. | https://www.graphpad.com/scientific-software/prism/ |
| WebLogo | (Crooks et al., 2004) | http://weblogo.berkeley.edu/logo.cgi |
| Geneious 11.0.2 | (Kearse et al., 2012) | https://www.geneious.com/ |
| R v 3.3.3 |  | https://www.r-project.org/ |
| Python v2.7 |  | https://www.python.org/download/releases/2.7/ |
| Rockhopper 2.0 | (Tjaden, 2015) | https://cs.wellesley.edu/~btjaden/Rockhopper/ |
| Edge-pro | (Magoc et al., 2013) | http://ccb.jhu.edu/software/EDGE-pro/index.shtml |
| DESeq | (Anders and Huber, 2010) | http://bioconductor.org/packages/release/bioc/html/DESeq.html |
| CLC Genomics Workbench 10 | (Qiagen) | https://www.qiagenbioinformatics.com/products/clc-genomics-workbench/ |
| MaxQuant | (Cox et al., 2014) | http://www.coxdocs.org/doku.php?id=maxquant:start |
| IterativeRank | This paper | https://github.com/mtien/IterativeRank |
| Sliding_window_analysis | This paper | https://github.com/mtien/Sliding_window_analysis |

## ACKNOWLEDGEMENTS

This project was supported by awards U19AI107792 (NIAID Functional Genomics Program) and 1R01GM087353 from the National Institutes of Health. We would like to thank members of the Crosson Lab for their contributions and input over the course of this project. The lab of Tao Pan lab provided important support in development of nucleic acid methods and lending equipment, most notably M.E. Evans and K.I. Zhou. Ruthenberg lab provided PP7 coat protein plasmids.

---

**Figure 1- Figure Supplement 1.**
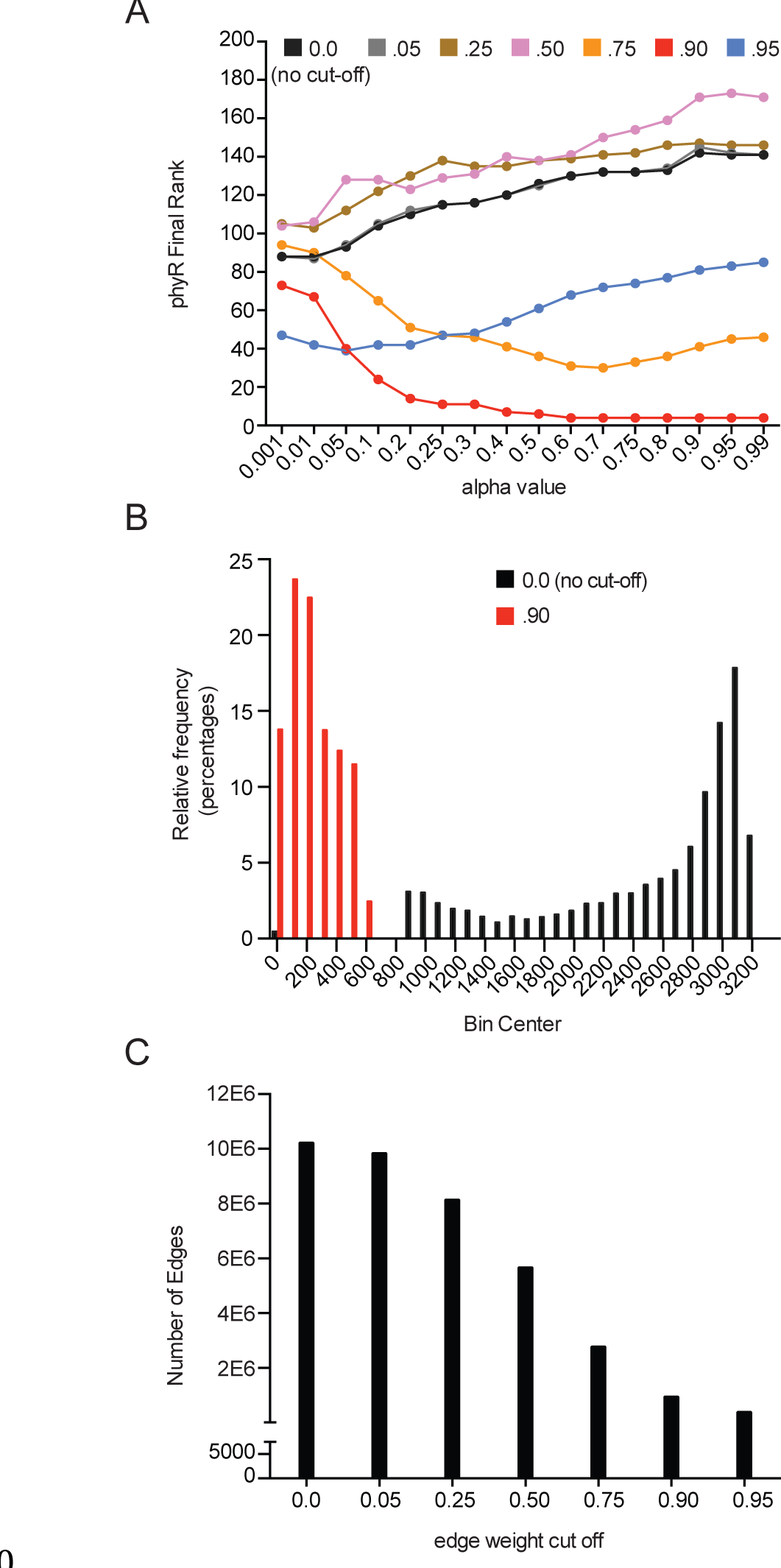
Parameter optimization of iterative rank through predicting *phyR* demonstrates edge-reduction as an important parameter. (see Materials and Methods - Iterative rank parameter tuning) (A) Systematic parameter exploration of the alpha value and edge-reduction, where *sigT*, *sigU*, *phyK*, *nepR*, *lovK*, and *lovR* are initialized with weight. Alpha value is plotted on the x-axis while the final rank of *phyR* on the y-axis. Colors indicate network where edges less than the indicated value were removed from the network. (B) The number of edges drawn for a given node shows that an edge reduction of 0.9 dramatically shifts the average number of edges per node. (C) The number of total edges in a network shows that an edge reduction of 0.9 is reduced by 10-fold.

**Figure 1- Figure Supplement 2.**
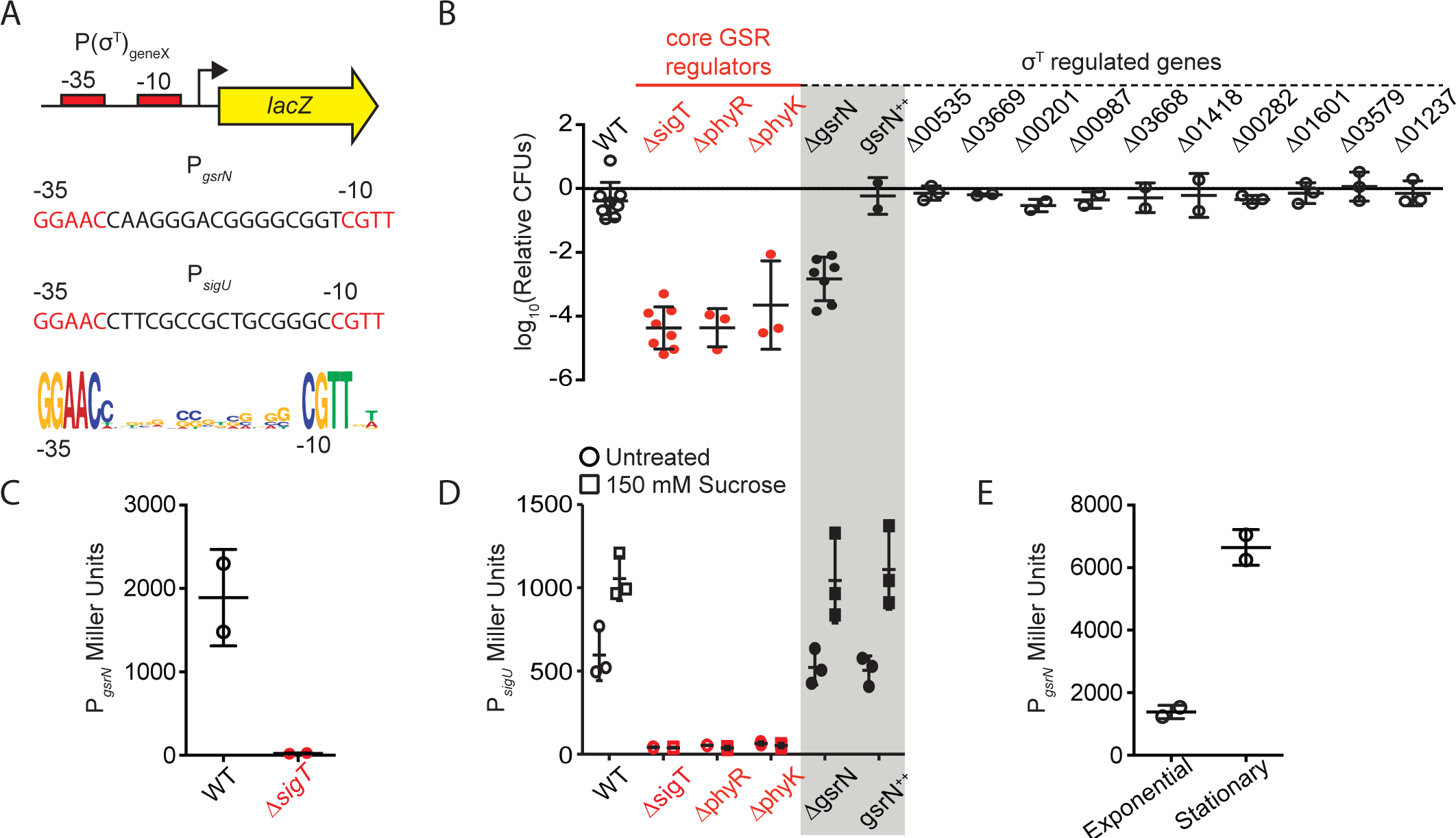
*gsrN* transcription is activated by σ^T^ and is induced under hyperosmotic stress and stationary phase growth. GsrN does not affect transcription from a σ^T^-dependent reporter. (A) Schematic of *lacZ* transcriptional fusions to the promoters of *gsrN* and *sigU. sigU* is a well-characterized reporter of GSR transcription (Foreman et al., 2012). Promoters of both genes contain a consensus σ^T^ binding site (nucleotides in red) (McGrath et al., 2007; Staron et al., 2009). σ^T^ binding motif (bottom) generated from twenty-one σ^T^-dependent promoters using WebLogo (Crooks et al., 2004). (B) Quantification of relative hydrogen peroxide survival of strains presented in Figure 1E. CFU of peroxide treated cultures were normalized by the CFU of the paired untreated culture. Genotypes are indicated above each bar. Numbers indicate CCNA locus numbers of deleted genes. *gsrN*^++^ is *gsrN* overexpression strain described in Figure 2A and Figure 2- figure supplement 1A. Bars represent mean ± SD from independent biological replicates (points). (C) β-galactosidase activity from the P*_gsrN_lacZ* transcriptional fusion in *Caulobacter* wild-type and Δ*sigT* backgrounds measured in Miller Units. Bars represent mean ± SD from 2 independent cultures. (D) β-galactosidase activity from the P*_sigU_lacZ* transcriptional fusion in a set of the genetic backgrounds in (B). GSR transcription was induced by exposure to 150 mM sucrose (final concentration) for three hours before measuring β-galactosidase activity. Bars represent mean ± SD from 3 independent cultures. (E) β-galactosidase activity from the P*_gsrN_lacZ* transcriptional fusion in exponentially growing (OD_660_ ^~^0.25) and stationary phase (OD_660_ ^~^0.75) wild-type cells. Bars represent mean ± SD from 2 independent cultures.

**Figure 2- Figure Supplement 1.**
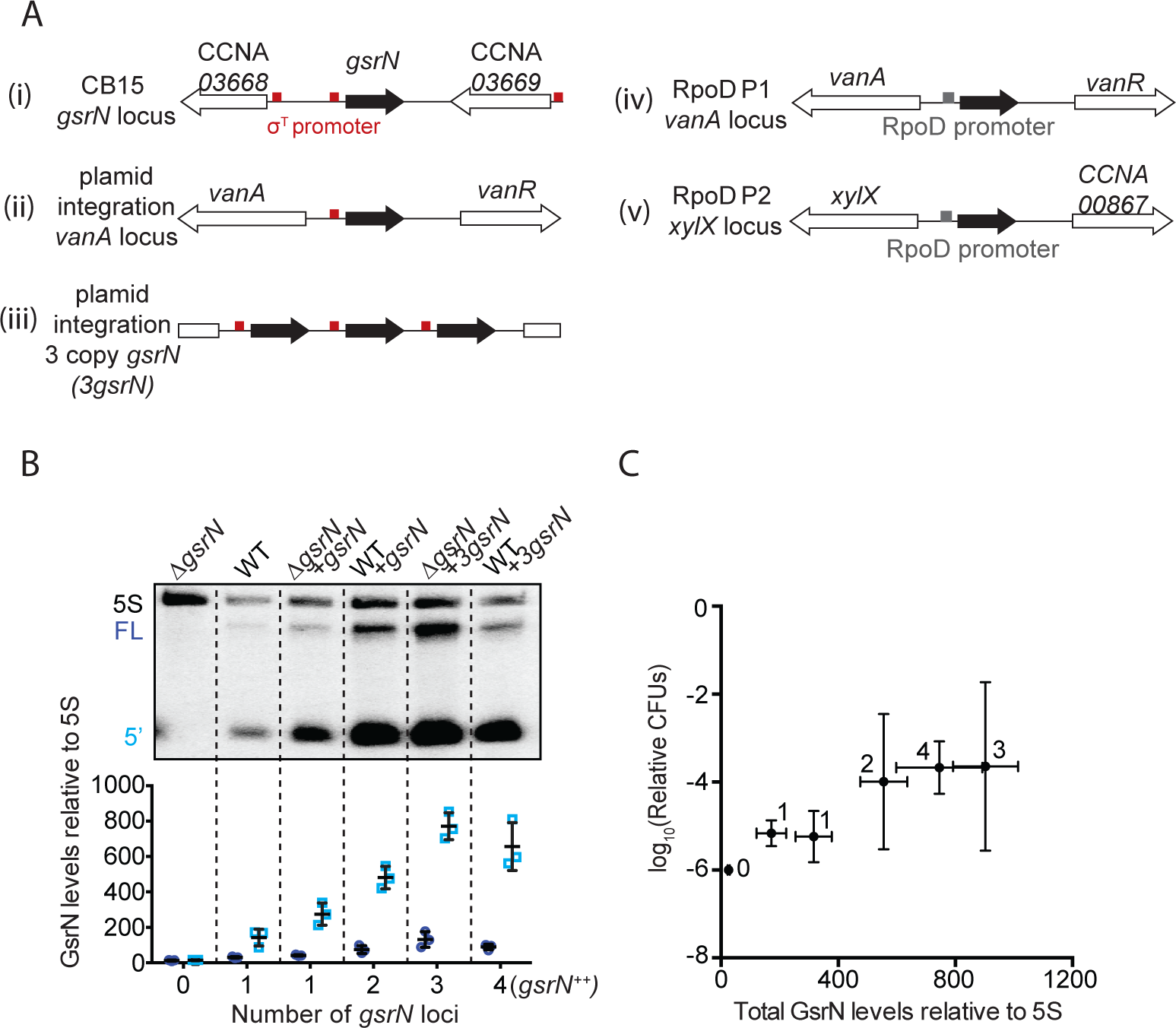
GsrN-dependent cell protection under oxidative stress is dose dependent. (A) In this study, *gsrN* was expressed several different ways. (i) At the native locus, the *gsrN* promoter contains a consensus σ^T^ binding site (red box); *gsrN* is flanked by two genes also with predicted σ^T^ promoters. Ectopic complementation and overexpression strains were created using pMT552-derived plasmids containing either (ii) one or (iii) three tandem copies of *gsrN* that were integrated into the chromosomal *vanA* locus. We also constructed strains in which *gsrN* expression was driven from one of two distinct σ^rpoD^-dependent promoters integrated in the chromosome. (iv) The RpoD1 promoter was taken from the predicted σ^rpoD^ binding site directly upstream of *vanA;* (v) RpoD2 promoter was taken from the predicted σ^rpoD^ binding site upstream of *xylX.* (B) Northern blots of RNA isolated from strains expressing increasing copies of *gsrN* probed with oligos complementary to GsrN. 5S rRNA was blotted as a loading control. Cells were harvested in exponential phase. Blots were quantified by densitometry. GsrN signal from the full-length (FL; dark blue) and 5’ isoform (5’; cyan) are normalized to 5S rRNA in each lane and multiplied by 100. Bars represent mean ± SD of triplicate extractions, each representing biologically independent samples. (C) Relationship between GsrN levels and peroxide stress survival. Total GsrN levels quantified by Northern blot (B) plotted against relative cell survival after 0.8 mM hydrogen peroxide treatment (CFU determined as outlined in Figure 1- figure supplement 2B). The Δ*gsrN* strain has zero CFUs after one hour treatment with 0.8 mM hydrogen peroxide, and no detectable GsrN by Northern blots, thus the y-axis point for this strain was plotted at 10^−6^, the detection limit of our assay.

**Figure 3- Figure Supplement 1.**
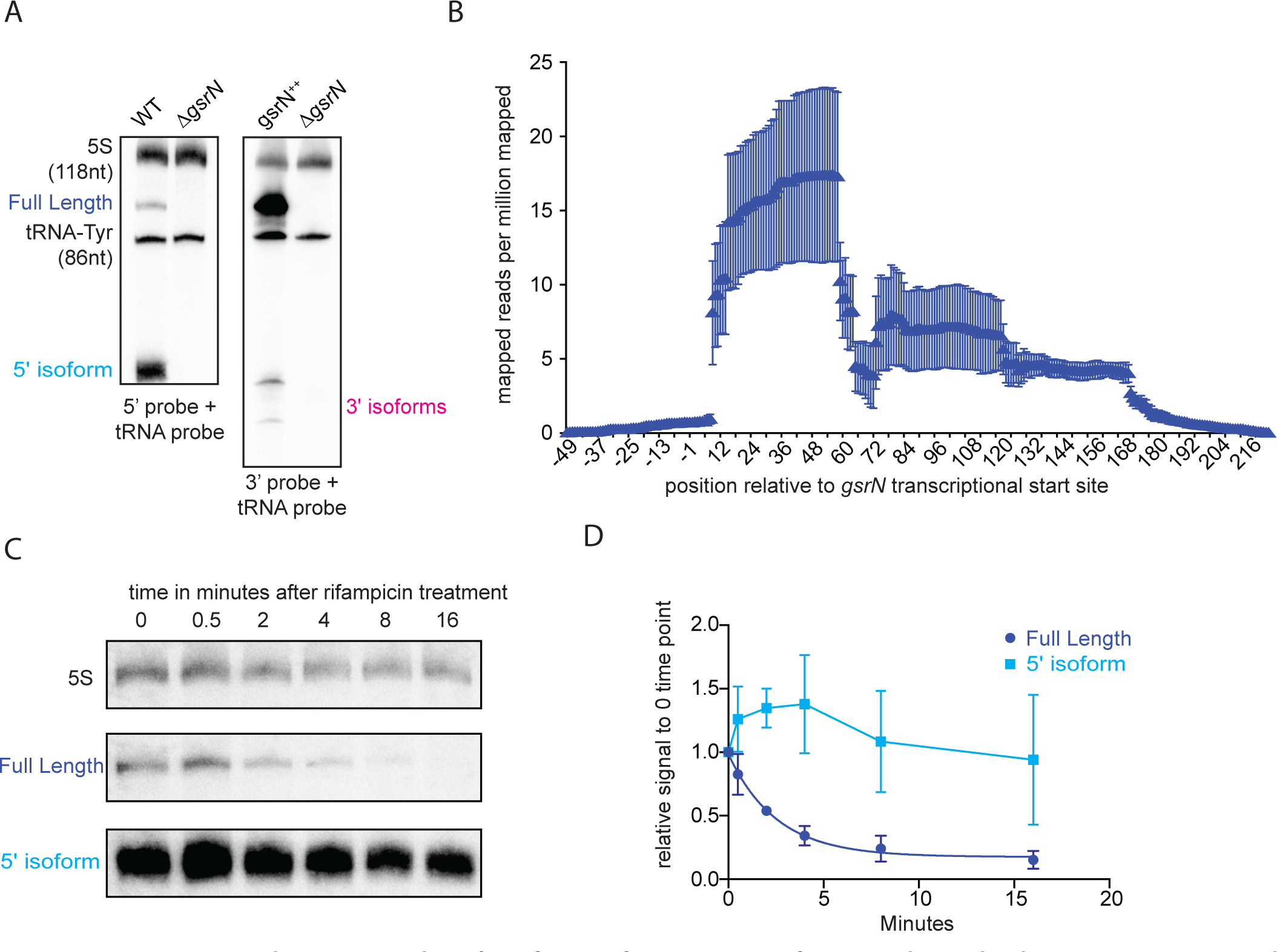
The 5’ isoform of GsrN arises from endonucleolytic processing and is the most abundant form of GsrN. (A) Northern blots of total RNA from cultures (OÜ660 ≈ 1.0) of wild type, Δ*gsrN*, and *gsrN^++^.* Blots were probed with ^32^P-labeled oligonucleotides complementary to either the 5’ or 3’ end of GsrN. Probes to 5S rRNA and tRNA-Tyr were used to estimate the size of full-length GsrN and its 5’ and 3’ isoforms. (B) RNA-seq read density from total wild-type RNA mapped to the *gsrN* locus. Chromosome position (x-axis) is marked in reference to the annotated transcriptional start site (TSS) of *gsrN* (position 3,830,130 in GenBank accession CP001340). Reads per million reads mapped is plotted as a function of nucleotide position. Mean ± SD from three independent biological replicates samples is plotted (GEO: GSR106168, read files: GSM2830946, GSM2830947, and GSM2830948). (C) Northern blot of total RNA extracted from wild type *Caulobacter* cells in exponential phase (OD_660_ ≈ 0.2-0.25) 0 to 16 minutes after treatment with 10 μg/mL rifampicin (final concentration). Bands for full-length GsrN, 5’ GsrN isoform, and 5S RNA loading control are shown. (D) Quantification of blots from (C) of full-length GsrN and 5’ GsrN isoform normalized to 5S rRNA levels in each lane. Signal at each time point is normalized relative to the zero minute time point. Data represent mean ± SD from three independent biological replicates.

**Figure 5- Figure Supplement 1.**
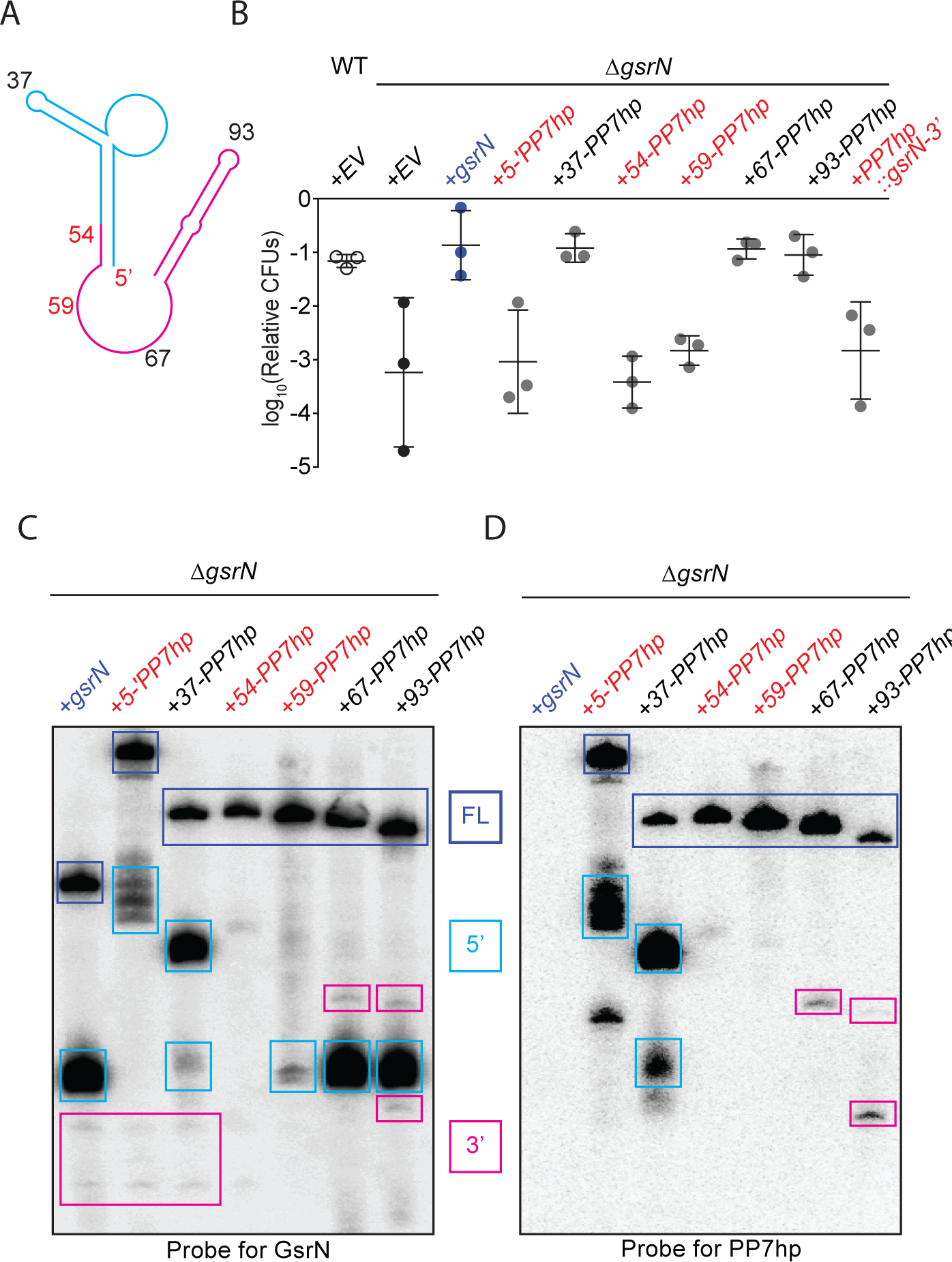
Identification, purification and biochemical characterization of functional GsrN-PP7hp chimeras. (A) Predicted GsrN secondary structure diagram from mFold (Zuker, 2003). Cyan and pink represent the 5’ and 3’ products, respectively, determined by primer extension and northern blot analyses (Figure 3). Numbered positions along the secondary structure indicate where PP7 RNA hairpin sequences (PP7hp) were inserted into *gsrN.* (B) Wild type, Δ*gsrN*-*EV*, and Δ*gsrN+gsrN*-*PP7hp* strains were subjected to hydrogen peroxide, diluted, and tittered as in Figure 1- figure supplement 2B. Empty vector (EV) strains carry pMT552. The nucleotide position of each PP7hp insertion in *gsrN* is marked above each bar. Data represent mean ± SD of three independent trials. (C) Northern blots of total RNA from stationary phase cultures (OD_660_ ≈ 1.0) of Δ*gsrN* strains carrying gsrN-PP7hp fusions. Blots were probed with oligonucleotides complementary to both the 5’ and 3’ ends of GsrN. Blot is overexposed to reveal minor products. Purple boxes mark full length GsrN, cyan boxes mark 5’ isoforms, and pink boxes mark 3’ isoforms. (D) Northern blots of same samples as in (C) ran in parallel but probed with oligonucleotides complementary to the PP7 hairpin sequence.

**Figure 6- Figure Supplement 1.**
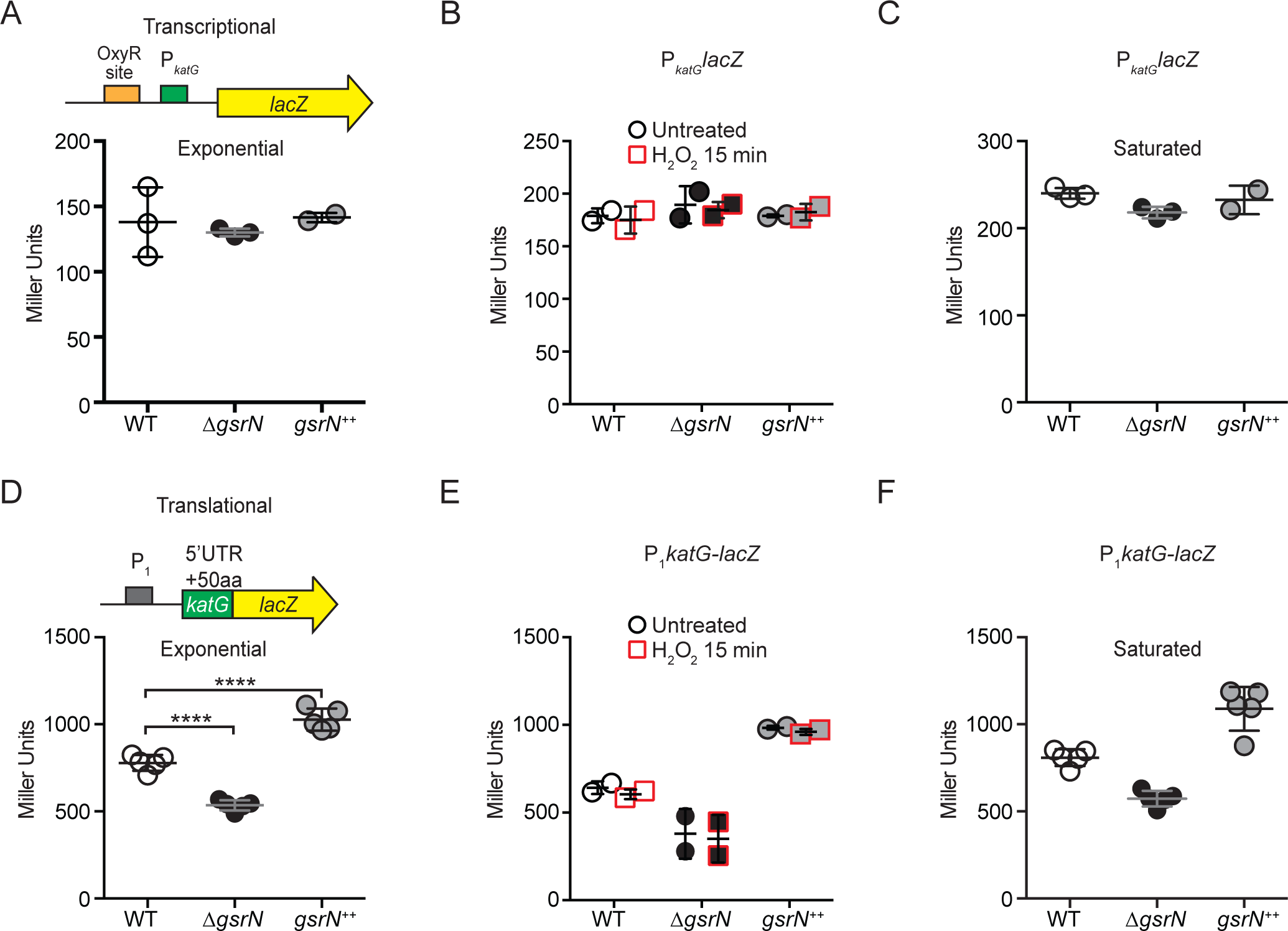
*gsrN* does not regulate *katG* transcription, but does enhance *katG*-*lacZ* mRNA translation. (A) *katG* transcriptional reporter construct contains the entire intergenic region upstream of *katG* fused to *lacZ* in pRKlac290. Transcription from this *katG* promoter (P_*katG*_) reporter was assayed in wild type, Δ*gsrN*, and *gsrN^++^* backgrounds during exponential growth (OD_660_ ≈ 0.2-0.25). Data represent mean ± SD of three independent trials. (B) Activity from the *katG* transcriptional reporter with and without a 15-minute treatment with 0.2 mM hydrogen peroxide. Cells were grown as in (A). Data represent mean ± SD of two independent trials. (C) Activity from the *katG* transcriptional reporter in stationary phase cultures (OD_660_ ≈ 1.0). Data represent mean ± SD of three independent trials. (D) KatG translational reporter (top) assayed in exponentially growing cells (bottom). Reporter is constitutively expressed from the P*_rpoD1_* promoter. *katG* leader (1-191 nt) region and the first 50 *katG* codons are fused in-frame to *lacZ.* Mean ± SD β-galactosidase activity, measured in Miller Units, presented from five independent trials. (E) Activity from the *katG* translational reporter was assayed in wild type, Δ*gsrN*, and *gsrN^++^* with and without a 15-minute treatment with 0.2 mM hydrogen peroxide. Cells were grown as in (A). Data represents mean ± SD from 2 independent trials. (F) Translation from *katG* leader fusion reporter was assayed in saturated cultures (OD_660_ ≈ 1.0). Data represents mean ± SD from 5 independent trials.

**Figure 6- Figure Supplement 2.**
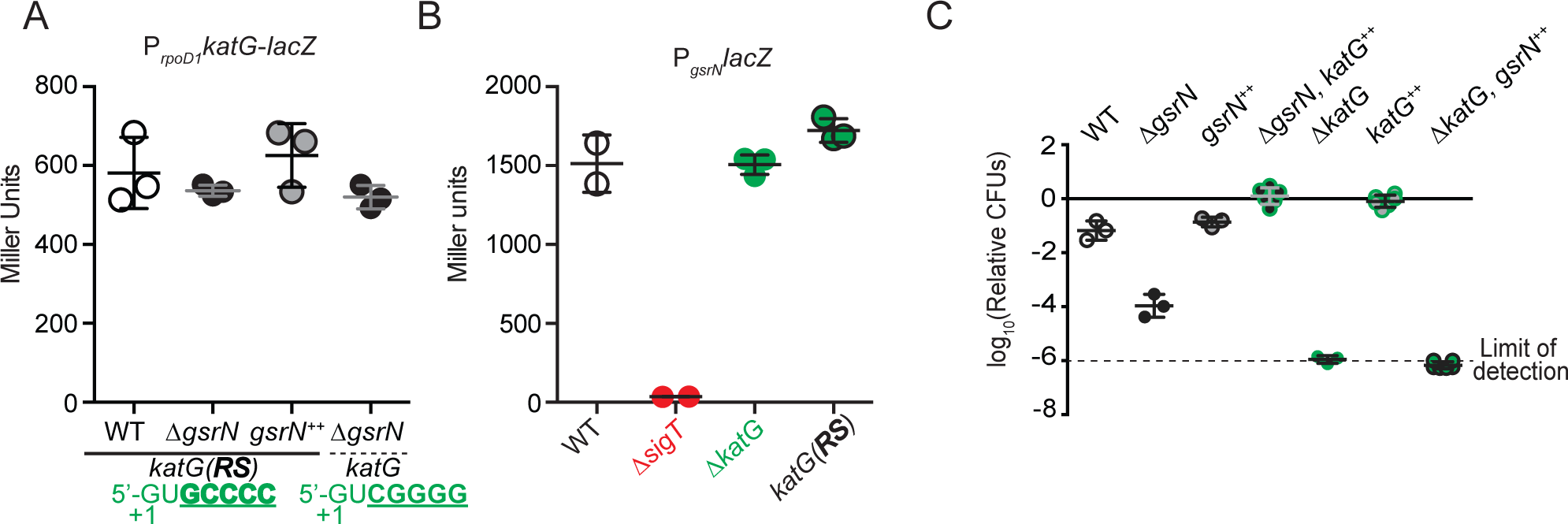
*katG(RS)-lacZ* translation is not affected by *gsrN. katG* does not affect *gsrN* transcription. *katG* is necessary and sufficient for peroxide stress survival. (A) Translational reporter activity from *katG* and *katG(RS)* leader fusions. The *katG(RS)*-*lacZ* construct is identical to that in (Figure 6- Figure Supplement 1D-F), except that it contains the reverse swapped target recognition site in the 5’ UTR upstream of *katG.* Activity from *katG(RS)*-*lacZ* in wild type, Δ*gsrN*, and *gsrN^++^* compared to *katG*-*lacZ* in Δ*gsrN* during exponential growth phase. Bars represent mean ± SD from 3 independent cultures. (B) Activity from the *gsrN* transcriptional reporter described in Figure S1A was assayed in *AkatG* and *katG-RS* backgrounds during exponential growth. Mean ± SD of 3 independent cultures. (C) Wild type, Δ*gsrN, gsrN^++^*, Δ*gsrN+katG^++^*(*P_xyl_katG*), Δ*katG*, *katG^++^*(*P_xyl_katG*), and Δ*katG*+*gsrN*^++^(4*gsrN*) strains were subjected to hydrogen peroxide, diluted, and tittered as in Figure 1- figure supplement 2B. Bars represent mean ± SD of at least 3 independent cultures.

**Figure 6- Figure Supplement 3.**
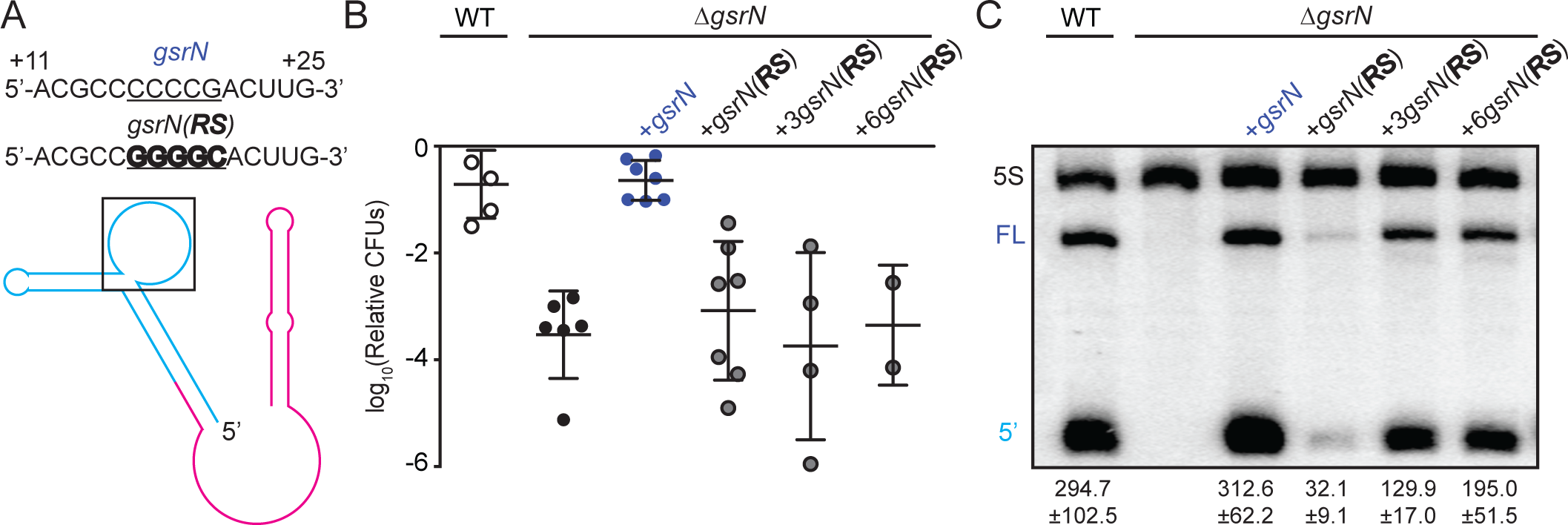
*gsrN* levels are determined by the sequence of its target recognition loop and *gsrN(RS)* allele cannot complement the peroxide susceptibility of Δ*gsrN*. (A) Predicted GsrN secondary structure diagram from Figure 3 with a black box is highlighting the nucleotides within the exposed 5’ loop of GsrN. Blue labeled sequence represents the wild-type *gsrN* coding sequence, with the underlined nucleotides emphasizing the location of the RS mutation. The bolded labeled sequence represents the RS mutant *gsrN* coding sequence. (B) Wild type, Δ*gsrN, ΔgsrN+gsrN* (complementation strain), and Δ*gsrN+XgsrN(RS)* strains were subjected to hydrogen peroxide, diluted, and tittered as in Figure 1- figure supplement 2B. Three different copy numbers of *gsrN(RS)* strains were tested. Bars represent mean ± SD of several independent cultures (points). (C) Northern blot of RNA extracted from wild type, Δ*gsrN*, and Δ*gsrN* complementation strains during exponential growth phase. Complementation strains include wild-type *gsrN* and reverse-swapped (RS) *gsrN(RS)* mutants. Three different copy numbers of *gsrN(RS)* strains were tested. Blots were probed with oligonucleotides complementary to the 5’ end of GsrN and to 5S rRNA. Quantified GsrN levels reported were normalized to the 5S rRNA signal in the same lane. Data represent mean ± SD from three independent biological replicates that were loaded, resolved, transferred, and hybridized on the same gel.

**Figure 8- figure supplement 1.**
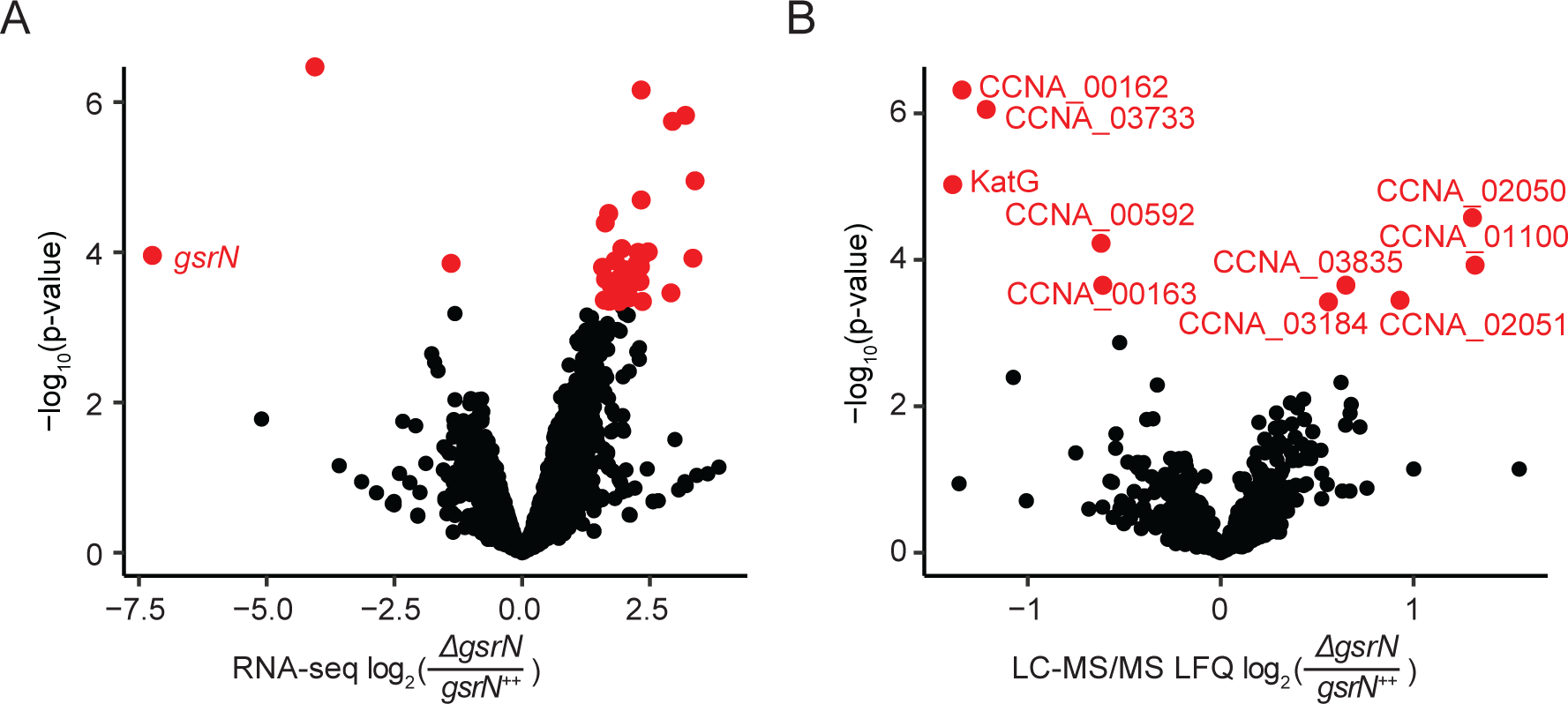
GsrN directly or indirectly affects the expression of multiple genes. (A) RNA-seq analysis of Δ*gsrN* and *gsrN^++^* early stationary phase cultures (OD_660_ ^~^0.85-0.90) represented as a volcano plot where expression changes are plotted as a function of p-value (Figure 8- source data 2). Red indicates transcripts with a false discovery rate (FDR) corrected p-value < 0.05. Black indicates gene transcripts with a FDR p-value above the cut-off. (B) LC-MS/MS total soluble protein signal from MaxQuant label free quantitation estimates from Δ*gsrN* and *gsrN^++^* cells grown to early stationary phase (OD_660_ ^~^0.85-0.90) (Figure 8- source data 3). Log-2 transformed fold change in LFQ estimates from MaxQuant (Cox et al., 2014) are plotted as a function of p-values obtained from the multiple t-test analyses using GraphPad Prism version 6.04 for MacOS, GraphPad Software, La Jolla California USA, www.graphpad.com. Red indicates proteins with significant differences (false discovery corrected p-value < 0.05). Black represents proteins that do not meet the FDR cut-off.

Gene Locus ID: GenBank locus ID Gene Name: if available

log_2_Fold: calculated fold change of the given region

Identification Method: refers to what strategy identified the enriched gene in the PP7hp affinity purification RNA-Seq

Region(s): the region and strand used to calculate the log_2_Fold metric. Additionally for the sliding window analysis additional information is provided. First letter indicates the relative position ofthe region indicated to the annotated gene coordinates. Briefly: I-internal, U-upstream, D-downstream. Second letter indicates the direction in which the reads mapped. Briefly: S-sense, A-anti-sense.

**Description: is the product description of the given gene(s)**

**Figure 1 - source data 1**

Excel file of gene expression data from (Fang et al., 2013) and estimated by Rockhopper (Tjaden, 2015). Each column represents the estimated expression from total RNA-extractions of *Caulobacter crescentus* cultures at five time points post-synchronization. These values were used to construct the network

**Figure 1 - source data 2**

Excel file of the results from the iterative rank algorithm. Results can be recapitulated using the scripts in https://github.com/mtien/IterativeRank.

**Figure 5 - source data 1**

Excel file of the output from Rockhopper analysis (Tjaden, 2015) on the RNA-Seq samples from the PP7 affinity purified total RNA samples. Figure 5C can be created using the python and R scripts in https://github.com/mtien/Sliding_window_analysis

**Figure 5 - source data 2**

Zipped file contain three files. These files include the sliding window analysis files generated from mapping the reads from the RNA-Seq experiment of the PP7 affinity purified total RNA samples. Figure 5D can be created using the scripts in https://github.com/mtien/Sliding_window_analysis

**Figure 5 - source data 3**

FASTA file that contains the windows of enrichment and total gene sequences of genes identified in the PP7 affinity purified total RNA samples.

**Figure 8 - source data 1**

Excel file that contains the log_2_Fold calculated values from both LC-MS/MS and RNA-Seq analysis of Δ*gsrN* versus *gsrN^++^*. Values used to calculate the fold changes from LC-MS/MS can be accessed from PRIDE: PXD008128, which contains the MaxQuant (Cox et al., 2014) LFQ protein group estimations under the name “MQrun_delta.txt” and “MQrun_plus.txt” representing the values for Δ*gsrN* versus *gsrN^++^*, respectively. Calculation of averages is outlined in Materials and Methods-LC-MS/MS processing of total soluble protein. Averages were then divided and log-transformed. Values used to estimate the fold changes from RNA-Seq were taken from the CLC workbench analysis of the GEO accession number GSE106168 files, see Materials and Methods-RNA-seq processing of total RNA.

**Figure 8 - source data 2**

Excel file that contains the compiled information from the CLC workbench analysis.

**Figure 8 - source data 3**

Excel file that contains the multiple t-test analysis outlined in Materials and Methods-LC-MS/MS processing of total soluble protein

